# Confirmation of preclinical findings across laboratories identifies robust effects and factors limiting reproducibility

**DOI:** 10.64898/2026.09.21.753112

**Authors:** Pasquale Pellegrini, Sophia C. Rotter, María Arroyo-Araujo, Clarissa F. D. Carneiro, Natascha I. Drude-Ratiu, Lorena Martinez Gamboa, Ana Banito, Nadine Bernhardt, Anne-Laure Boulesteix, Frank Buttgereit, Astrid Dempfle, Nayar Durán-Hernández, Timo Gaber, Marcus Groettrup, Bettina Habelt, Bernhard Haller, Hazem Hamza, Simone Hettmer, Anca Kliesow Remes, Sebastian Kobold, Merle Kochan, Frank Konietschke, Lorenz H. Lehmann, Max Löhning, Andrea Matzen, Marcus W. Meinhardt, Helen Morrison, Oliver J. Müller, Oliver Planz, Günther H. S. Richter, Lars B. Riecken, Steffen K. Rosahl, Kerstin Rubarth, Katharina Schmidt-Bleek, Lena Schuler, Arunabh Sharma, Ivan Skorodumov, Rainer Spanagel, Irene Teichert-von Luettichau, Matthias Tenbusch, Mohsen Valadan, Juliane C. Wilcke, Ulf Toelch

**Author notes:** These authors contributed equally to this work (Pasquale Pellegrini, Sophia C. Rotter). Instituto Serrapilheira, Rio de Janeiro, Brazil (Clarissa F. D. Carneiro). Department of Medicine V, University Medical Centre Mannheim, University of Heidelberg, Mannheim, Germany (Lars B. Riecken). This author passed away prior to publication (Marcus Groettrup).

## Abstract

A fundamental challenge in drug development is the frequent failure of early laboratory research to translate into clinical benefit. One proposed solution is to confirm findings from exploratory single-laboratory studies across multiple laboratories before clinical testing. We investigated this approach by prospectively following the conduct of twelve preclinical multi-laboratory studies across different fields of medicine and by systematically identifying previously published multi-laboratory studies in the literature. To explore which factors determine confirmation success, we evaluated effect sizes, experimental rigor, and a set of predefined criteria. When tested under increased rigor, only a fraction of multi-laboratory studies confirmed their initial results. Effect size shrinkage between exploratory and confirmatory stages was attributable to shifted control baselines, weaker treatment responses, or a greater variance inflation. In summary, multi-laboratory studies proved complex yet highly informative and serve as an effective filter for preclinical treatment candidates.

## Introduction

A fundamental challenge in biomedical research is the frequent failure of preclinical evidence to translate into clinical benefit. Currently, the pharmaceutical pipeline is time-consuming with high attrition rates (*1*). New interventions show promise in preclinical efficacy studies, but clinical trials ultimately fail to reproduce the same effect in over 90 % of cases (*2–6*). Even though the causes of these failures are multifactorial, lack of preclinical predictivity, meaning the extent to which preclinical results correlate with clinical benefit in patients, is recognized as a principal determinant (*7*). For preclinical findings to be predictive, they must first be reproducible and generalizable (*7*). When evaluating the reproducibility of early preclinical findings, retrospective investigations have found low success rates (*8–11*).

One reason is that current preclinical approaches are primarily exploratory, frequently lacking two key aspects: reliability and validity (*12*). Reliability refers to the consistency of results under repeated testing. Larger, adequately powered studies produce more precise effect estimates with narrower confidence intervals and are therefore more likely to yield reproducible results (*13*, *14*). Validity refers to whether an experimental approach measures what it intends to measure. It is improved by implementing strategies such as randomization and blinding to mitigate risks of bias (internal validity) (*15*), by broadening the generalizability of findings (external validity) (*16*), and by incorporating clinically relevant features (translational validity) (*17*). Therefore, initial results from exploratory research require follow-up studies – *confirmatory studies* – that improve validity and reliability (*17–20*). Whereas there is a clear objective to use these studies as a critical step toward clinical trials, the exact methodological setup and concrete implementation remain scarce and understudied.

Here, we investigated two datasets of preclinical confirmatory studies that tested the significant results of an initial single-laboratory (single-lab) exploratory study in a rigorous, multi-laboratory (multi-lab), and translation-oriented setting. The *primary confirmatory studies (pCS)*, the first set, stem from an initiative of the German Federal Ministry of Research, Technology and Space (BMFTR), which funded twelve such studies in 2020 (*21*). For this dataset, we had access to in-depth information on grant proposals and partly unpublished exploratory and confirmatory study designs and results. This offers a unique perspective on the research process for a complete sample of preclinical confirmatory studies conducted under similar conditions. This dataset is complemented by a second set, the *extended confirmatory studies (eCS),* which is a collection of published multi-lab studies. These are, like the *pCS*, each based on a specific single-lab exploratory study (Fig. 1A, tabs. S1 and S2), and provide a published sample of similar studies that do not originate from a common funding line. Importantly, both datasets differ from recent initiatives that conducted direct replications, repeating prior experiments using the same methods or models (*10*, *11*). Replications aim to determine whether the same experimental results in a specific field (e.g., oncology (*10*)) or country (e.g., Brazil (*11*)) can be obtained again under the same or very similar conditions. Confirmatory studies extend replications by incorporating systematic changes and extensions regarding validity and reliability. This includes, for example, blinding and randomizing, testing both sexes, increasing sample sizes, or testing in a multi-lab setup. Our two unique datasets allow a detailed assessment of how preclinical confirmatory studies may identify potential treatment candidates and index those that potentially fail downstream. We investigated how outcomes changed between exploratory and confirmatory stages and deconstructed how these changes emerged. Specifically, we applied different criteria to assess confirmation success and examined methodological differences between stages. Through this, we mapped how preclinical confirmatory studies provide useful diagnostic information about underlying knowledge claims and inform future experimental steps.

**Fig. 1.**
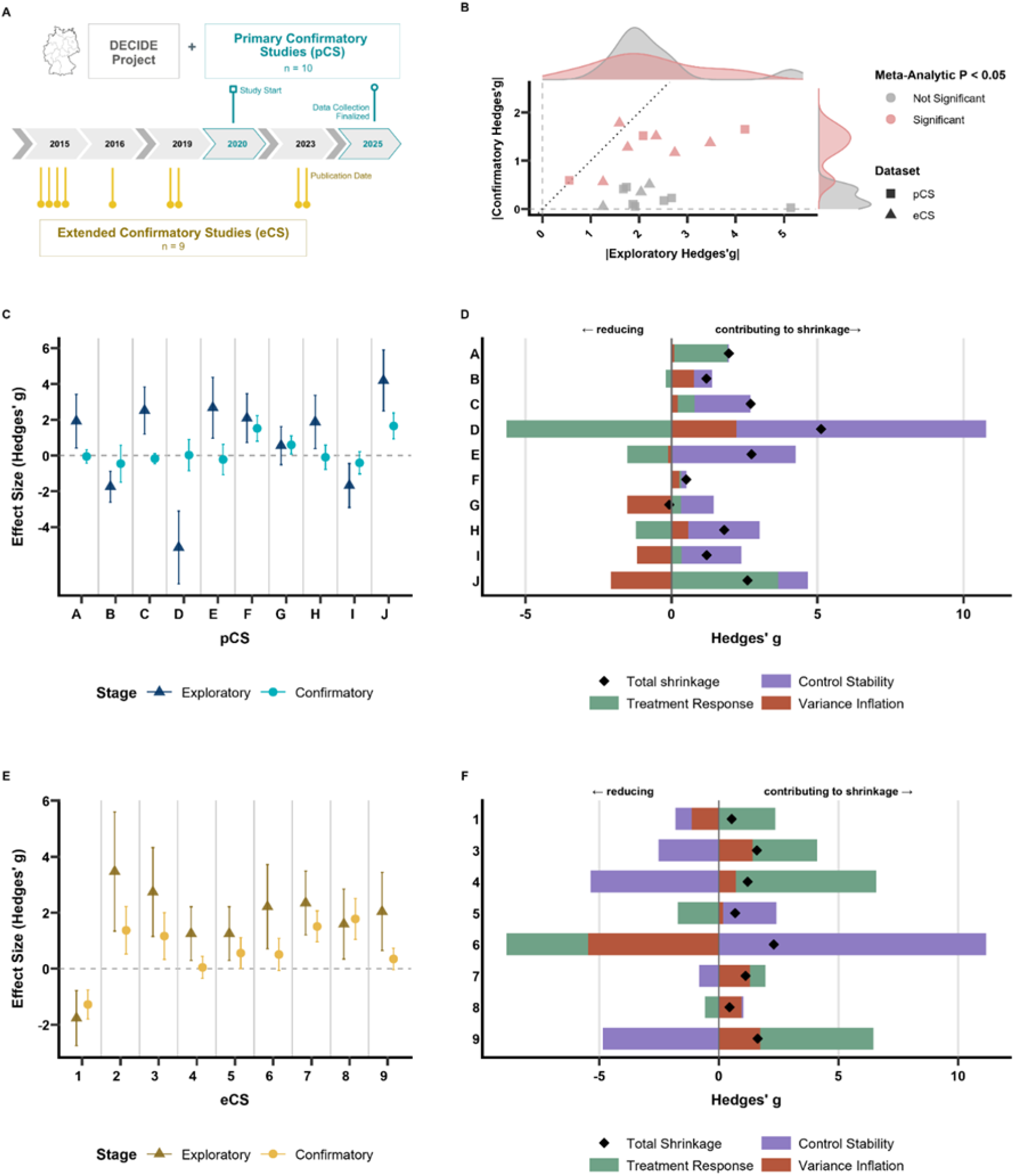
Effect size attenuation and less statistically significant results in multi-laboratory (multi-lab) studies. (A) Overview and timeline of the primary confirmatory studies (pCS), multi-lab studies conducted with DECIDE team support, and publication dates of the extended confirmatory studies (eCS), published multi-lab studies identified through a systematic literature search. (B) Exploratory versus pooled multi-lab confirmatory effect sizes (Hedges’ g) for pCS and eCS; each point is one study. The dashed line is the identity line (no shrinkage). Red: statistically significant pooled confirmatory effects (fixed-effects model, P < 0.05). Meta-regression slopes (tab. S5): eCS 0.44 (95% CI 0.22 to 0.66, P < 0.001); pCS 0.14 (−0.03 to 0.31, P = 0.11); difference −0.30 (−0.58 to −0.02, P = 0.034). (C and E) Exploratory (single-lab) and confirmatory (pooled, fixed-effects) effect sizes for pCS (C) and eCS (E). Error bars: 95% confidence intervals. (D and F) Decomposition of shrinkage from exploratory to confirmatory stage for pCS (D) and eCS (F). Bars show the contribution of control stability, treatment response and variance inflation to the change in Hedges’ g per study (A–J, pCS; 1–9, eCS); positive values contribute to shrinkage, negative values counteract it. Black diamonds: total observed shrinkage.

## Results

### Systematic experimental differences drive effect size shrinkage

Our analysis comprised ten primary confirmatory studies (*pCS*), defined as multi-lab studies with a corresponding single-lab exploratory study. Of the twelve studies initially funded, two were discontinued prematurely due to technical or regulatory reasons. The extended confirmatory dataset (*eCS*), identified through a systematic literature search (tabs. S3 and S4), provided nine additional pairs of exploratory and confirmatory studies for comparison. Individual studies are identified throughout by letters (*pCS:* A–J) and numbers (*eCS:* 1–9). We first assessed the success of confirmatory studies using effect size and statistical significance as the primary metrics. Exploratory studies across both datasets exhibited medium to large effect sizes (Hedges’ *g*pCS = 0.73 ± 2.79; *g*eCS = 1.68 ± 1.48; mean ± SD), but subsequent confirmatory multi-lab studies showed substantial effect size shrinkage (Fig. 1B). The effect size reduction was stronger in the *pCS* than in the *eCS* (tab. S5). Based on a fixed-effect meta-analysis, three of ten *pCS* (30%) and six of nine *eCS* (67%) confirmed the exploratory results (Fig. 1, C and E; tabs. S6 and S7). Since all confirmatory results were based on an exploratory finding, we deconstructed the effect size shrinkage into three factors attributable to differences between experimental stages: *control instability* (the control baseline shift from exploratory to confirmatory stage), *treatment response attenuation* (a reduction in the mean of the treated group), and *variance inflation* (the increase in primary outcome variability within and between labs) (Fig. 1, D and F; see Materials and Methods).

Effect size shrinkage in both *pCS* and *eCS* was differentially driven by these three factors across individual studies. The *pCS* demonstrated large overall shrinkage in effect size magnitude (0.51 – 5.12 Hedges’ *g*), stemming from a mix of treatment response attenuation, control instability, and variance inflation (Fig. 1D). In project G, opposing factors neutralized the net shrinkage. *eCS* shrinkage ranged from moderate to very large (0.45 – 2.3 Hedges’ *g*), reflecting a heterogeneous contribution of all three factors across projects (Fig. 1F). Effect size shrinkage potentially arises from differences between labs. We therefore assessed inter-laboratory (inter-lab) variability using the distance method (*22*), which quantifies how far each lab’s observed mean difference deviates from the overall multi-lab mean, relative to the total variability across all labs. Labs exceeding chance-level deviation were flagged as atypical (Fig. 2, A and B). No atypical labs were detected in the *pCS*, possibly reflecting low test sensitivity in two-lab studies or small inter-lab variance (fig. S1, A and B). In the *eCS*, studies four and six each contained one atypical lab (Fig. 2B). Study six illustrates that with only two labs, results can diverge strongly, yet it is often impossible to determine which laboratory reflects the true effect and which deviates from it. Overall, inter-lab variability was comparable across the two datasets. That is, there were no substantial differences in either absolute (Root Mean Squared Error, RMSE) or relative (Coefficient of Variation, CV) inter-lab variability (Wilcoxon rank-sum test; RMSE: *W* = 28, *n*₁ = 10, *n*₂ = 9, *P* = 0.182; Fig. 2C; CV: *W* = 54, *n*₁ = 10, *n*₂ = 9, *P* = 0.49; fig. S1C).

**Fig. 2.**
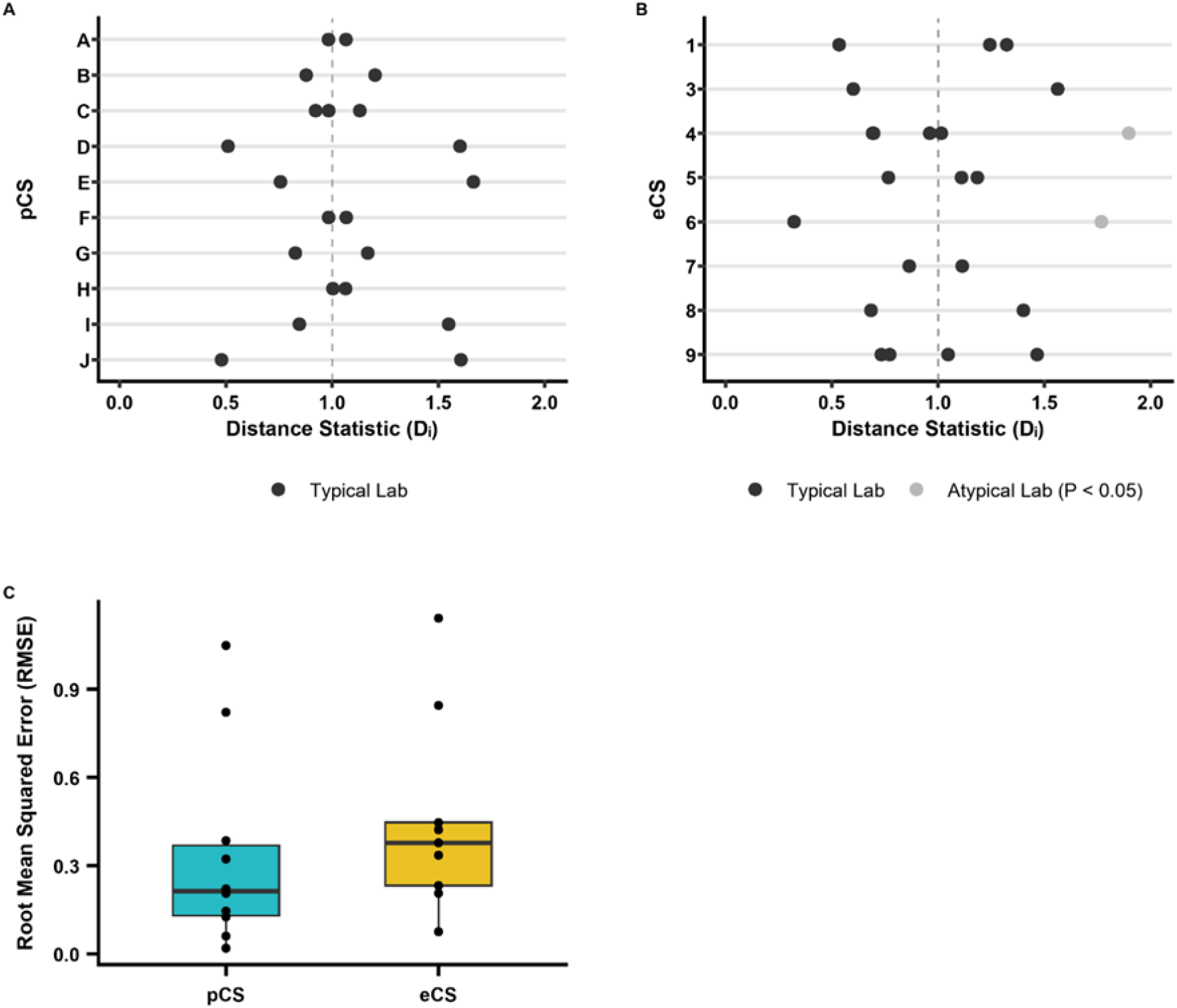
Inter-laboratory variability in confirmatory studies. **(A** and **B)** Distance Dᵢ (mean-difference *F*-statistic) for *pCS* (A) and *eCS* (B): quantifies how different each laboratory’s mean difference is relative to the grand mean across laboratories. Values exceeding the critical threshold (grey dots) indicate statistically atypical laboratories (*P* < 0.05). The dashed line at Dᵢ = 1 indicates no deviation from the expected distance based on the grand mean. **(C)** Inter-laboratory variability in effect size estimates, quantified as the root mean square error (RMSE) of Hedges’ *g* relative to the pooled multi-lab estimate, shown for *pCS* and *eCS*. Boxes indicate the median and interquartile range (IQR); whiskers extend to 1.5 × IQR, and points represent individual studies. Group differences were assessed using a two-sided pairwise Wilcoxon rank-sum test with Benjamini-Hochberg correction (*n*pCS = 10, *n*eCS = 9, *W* = 28, *P* = 0.182).

### Confirmation success remains low across alternative criteria

Observed results contrasted starkly with the *Expected Replication Rate* (ERR), a z-curve-based estimate of replicability from the distribution of significant findings (Fig. 3A). Based on the exploratory studies, the analysis indicated a joint 85% expected replication rate compared to an average confirmation success of 47% across *pCS* and *eCS (fixed-effect models,* tabs. S6 and S7). ERR assumes a strict significance threshold for confirmatory testing. Alternative criteria to assess the congruence of exploratory and confirmatory results exist (Fig. 3B, tab. S8). We, therefore, tested how a set of criteria evaluated confirmatory studies for success. For this, we ran a simulation to test the accuracy of these criteria in a scenario reflecting the empirical exploratory-to-confirmatory setup. Our model accounted for the observed effect size shrinkage, whereby strong initial findings become weaker in later, stricter tests. This allowed us to estimate how well each criterion identified true success, despite shrinkage. We also included a scenario with complete shrinkage to measure how often our criteria produced false positives (see Materials and Methods).

**Fig. 3.**
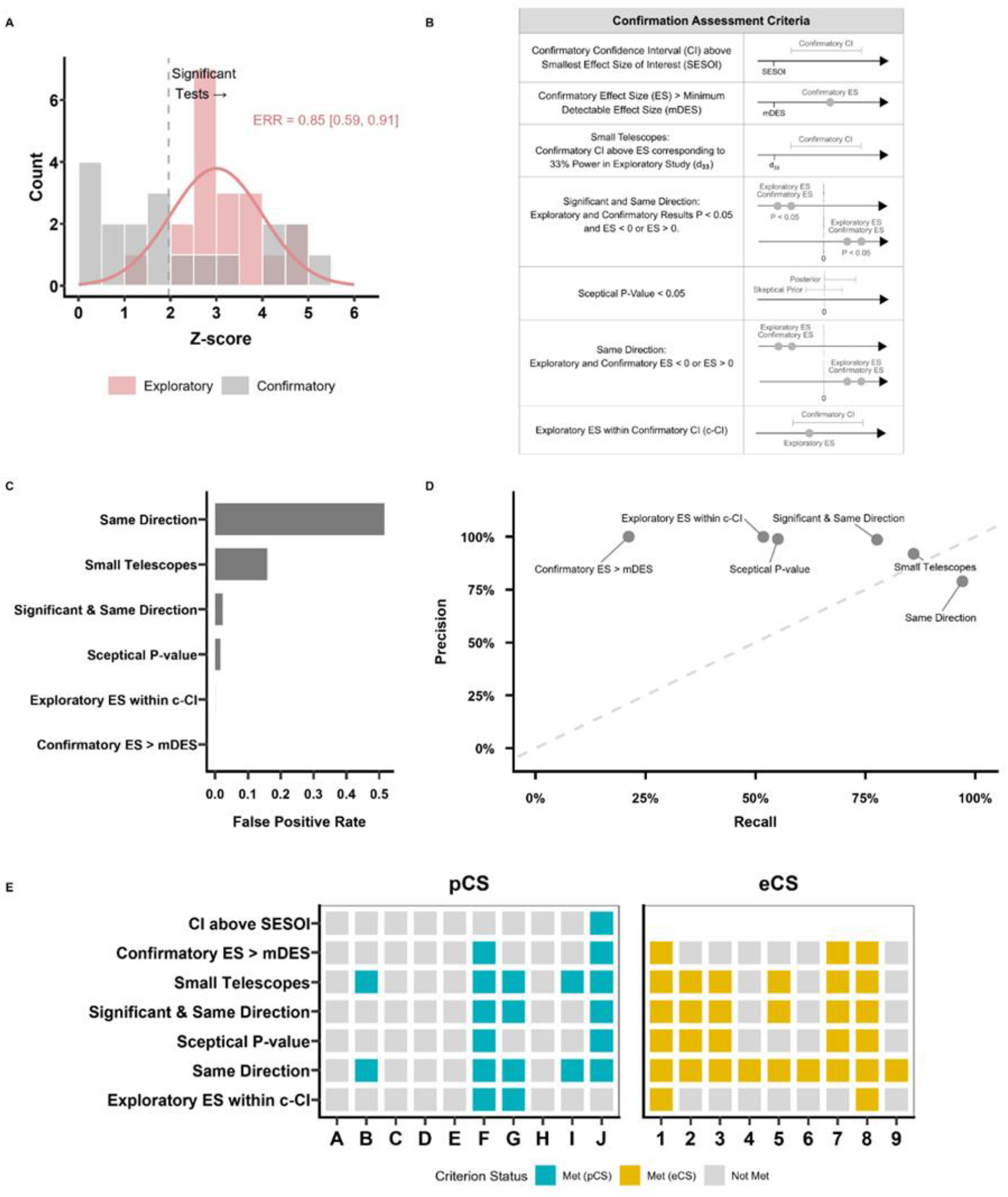
Expectation and confirmation success across multiple decision criteria. **(A)** Z-curve analysis of statistically significant test statistics (*pCS* and *eCS* combined), converted from two-sided *P*-values (capped at *z* = 6). The dashed vertical line indicates the significance threshold (*z* = 1.96). The Expected Replication Rate (ERR), the model-estimated average true power among significant exploratory findings, was estimated from the exploratory distribution. Confirmatory values are shown for comparison only. **(B)** Overview of the seven criteria and their decision rules (criteria in the Supplementary Materials). **(C)** False positive rate (FPR) for each criterion under the null hypothesis (99% shrinkage). Bars show simulated trajectories misclassified as successful despite a near-null confirmatory effect, among those passing the exploratory threshold (*P* < 0.05). “CI above SESOI” was excluded, as it requires an empirically defined SESOI unavailable for simulated trajectories. **(D)** Precision-recall plot for the decision criteria. True positives were defined as 0% or 20% shrinkage, and true negatives as 99% shrinkage. Each point represents one criterion. **(E)** Confirmation success matrix for *pCS* (left) and *eCS* (right) evaluated against the indicated criteria (rows). Colored cells indicate a criterion was met. Abbreviations: SESOI, smallest effect size of interest; mDES, minimum detectable effect size; c-CI, confirmatory confidence interval.

Based on this assessment (Fig. 3, C and D), several criteria classified a higher proportion of studies as successful compared to the standard P-value-based criterion (Fig. 3E). However, even the most lenient criterion (“Same Direction”, Fig. 3D) indicated only a 74% success rate across *pCS* and *eCS*. Such lenient criteria also showed elevated false positive levels (Fig. 3C). Stricter criteria, such as the “Sceptical P-Value”, identified only 36% of all studies as successful, which was similar to the “Significant and Same Direction” criterion. Successful confirmation thus remained low regardless of the applied criterion.

### Confirmatory multi-lab studies apply high methodological standards

Confirmatory studies showed an increase in validity and reliability relative to exploratory experiments. Although per-lab sample size was unchanged between stages (fig. S2, tab. S9), *pCS* and *eCS* achieved 2.8-fold larger overall sample sizes across labs on average, reflecting the participation of additional labs at the confirmatory stage. This addressed a key limitation of exploratory research, namely that small sample sizes may fail to detect true effects (Type II error). Consequently, confirmatory studies could detect smaller effects compared to exploratory studies (Bayesian Lognormal Hierarchical Regression: β = -0.62, 95% CrI [-0.87, -0.36]; Fig. 4A, tab. S10), considerably increasing reliability. This indicates that the low confirmation rates above are unlikely due to insufficient statistical power at the confirmatory stage.

**Fig. 4.**
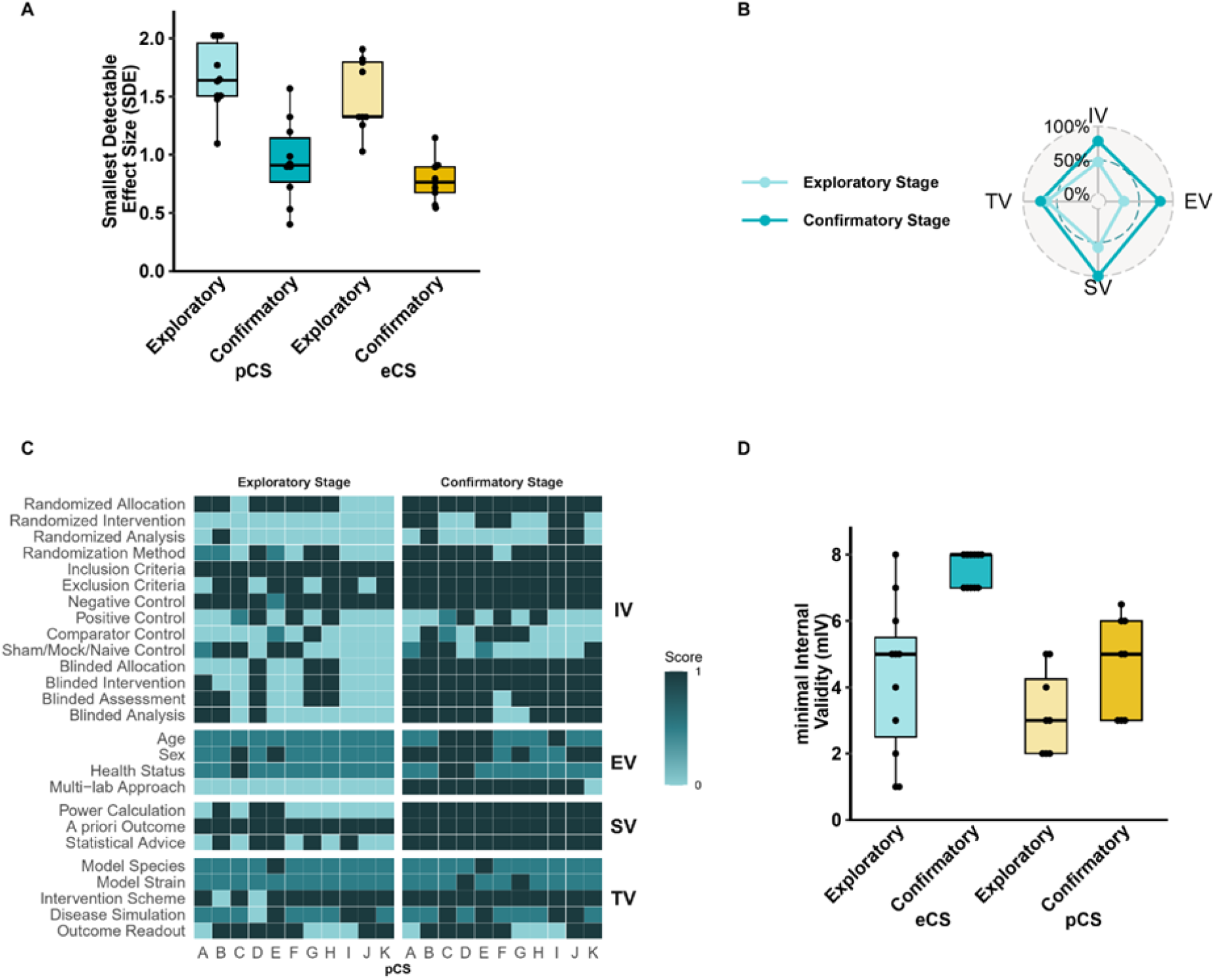
Increased methodological rigor in preclinical confirmatory studies. **(A)** Smallest detectable effect size (SDE) in the exploratory and confirmatory stages for *pCS* and *eCS*. SDE is the minimum effect size detectable with 80% power at α = 0.05 given each study’s total (pooled) sample size, not accounting for within-laboratory correlation. Assessed with a Bayesian hierarchical lognormal regression (stage × dataset interaction, random intercept for study); credible intervals excluding zero indicate a credible difference. **(B)** Total scores across the four validity domains (IV, internal; EV, external; SV, statistical; TV, translational validity) for *pCS* (*n* = 11) at exploratory and confirmatory stages, rescaled to 0–1 and averaged by stage and domain. **(C)** Heatmap of validity scores for individual validity items (left labels; e.g., blinding, randomization) and validity domain (right labels; IV, EV, SV, TV) across exploratory and confirmatory stages (*n* = 11). **(D)** Comparison of minimal internal validity (mIV) scores between *pCS* and *eCS*. Assessed with a Bayesian hierarchical ordinal (cumulative probit) regression (stage × dataset interaction, random intercept for study). Credible intervals excluding zero indicate a credible difference. One *eCS* data point was excluded to avoid double-counting a study with two outcomes.

For the *pCS*, we obtained in-depth information on the experimental design through a survey of the studies’ researchers. Based on this survey, we developed a scoring system to quantify changes in experimental rigor between the exploratory and confirmatory studies (tab. S11). Compared to single-lab exploratory experiments, *pCS* showed substantially higher internal, external, and statistical validity (Fig. 4B). To increase internal validity, each *pCS* implemented blinding for group allocation and intervention administration; most also blinded outcome assessment and analysis (Fig. 4C). All *pCS* randomly allocated subjects into groups, often via computer-generated block randomization accounting for lab, sex, weight, or age. Each study defined *a priori* inclusion and exclusion criteria to limit selection bias. To increase external validity, *pCS* harmonized protocols across labs and often included both sexes and varying ages (Fig. 4C). All *pCS* performed a *priori* sample size calculations to achieve adequate power and were advised by a biostatistician, increasing statistical validity (Fig. 4C). Translational validity was addressed via model- and intervention-specific strategies. Study B, for example, shifted from a prophylactic to a therapeutic intervention scheme, whereas others adopted animal strains, species, or model inductions with increased translational relevance (Fig. 4C). Consequently, the four validity domains (internal, external, statistical, translational) separated confirmatory from exploratory studies in an unsupervised clustering approach (fig. S3).

Whereas both datasets shared many characteristics (tab. S1), survey-level detail on experimental design was available only for the *pCS*, not the *eCS*. We therefore compared studies using a preregistered minimal internal validity (mIV) score assessing randomization, blinding, controls, and population criteria extracted directly from the *eCS* publications (tab. S12). Overall, confirmatory studies showed higher mIV scores than their corresponding exploratory studies (Bayesian Ordinal Hierarchical Regression: β = 2.55, 95% CrI [1.52, 3.69]). For the *eCS*, however, this increase was attenuated and markedly lower than for the *pCS* (β = -1.54, 95% CrI [-2.86, -0.25]) (Fig. 4D, tab. S13).

## Discussion

Given the low reproducibility of preclinical findings and the frequent failure of their clinical translation, new approaches are needed to increase the efficiency of preclinical research. Here, we investigated how confirmatory multi-laboratory designs may increase the efficiency of preclinical research by facilitating the identification of potential treatments, using two independent datasets of preclinical studies. Confirmatory studies were conducted with higher rigor but exhibited lower effect sizes than exploratory studies. As a result, only a subset of studies confirmed their initial results. Inflated effect sizes in exploratory studies typically arise from a phenomenon called the winner’s curse (*23*). It occurs in underpowered experiments with small sample sizes (*13*, *14*) and is most prevalent in early discovery phases. Here, unrepresentative samples frequently and systematically overestimate the true size of an effect. Consequently, effect sizes frequently shrink under more rigorous confirmatory testing (*10*). A standardized effect size expresses the difference between treatment and control group means relative to the variability of the data. It can therefore shrink for two reasons: the difference between the groups becomes smaller, or the variability within the groups becomes larger. In our sample, the observed shrinkage could be traced back to specific differences between exploratory and confirmatory stages. In some studies, control group means differed substantially between stages. If animals of the negative control are healthier at the confirmatory stage, a reduced effect size does not necessarily mean the intervention lacks therapeutic efficacy. Some confirmatory studies showed an increased treatment variance, a well-documented phenomenon in human clinical trials (*24*). This heterogeneity may reflect variable individual response to treatment, as non- or low-responders inflate the variance and thus reduce the overall effect size. In such cases, exploratory subgroup analyses are crucial for informing subsequent experimental designs. Establishing *a priori* minimum responder thresholds, similar to clinical trials, potentially mitigates this effect, qualifying it as an additional strategy for preclinical confirmatory studies (*25*).

Effect size shrinkage can also reflect differences between labs. Some projects showed considerable inter-lab variability, resulting in lower overall effect sizes through outcome-relevant lab idiosyncrasies such as experimenter effects (*26*). To avoid this, multi-lab studies frequently harmonized their protocols, thereby reducing variance between labs, particularly in the *pCS*. A recent preclinical stroke study similarly harmonized operating procedures across labs through joint workshops, greatly reducing the coefficient of variation in the primary outcome and resulting in comparable means across labs (*27*). Harmonization thus reduces random, unintended inter-lab variability.

Confirmatory studies can effectively inform decisions about next research steps. To assess how sensitive such decisions are, we tested confirmation success against several criteria. An *in silico* investigation of the test stringency across different criteria revealed differential balancing of diagnostic measures for each criterion. “Significant and Same Direction” and the “Sceptical P-Value” (*28*) represent conservative criteria with high positive predictive values, consistent with a recent simulation study (*29*). Other criteria, such as “Small Telescopes” (*30*) and “Same Direction”, were more liberal, with higher false positive rates. This highlights the importance of upfront statistical planning and choosing a criterion that serves the desired goal: either to eliminate ineffective candidates or to avoid missing efficacious ones. In conjunction with the analysis of the effect size shrinkage, this gives a nuanced view of confirmatory study results that would not be possible through single-lab studies alone. We classified several studies as successes under conservative criteria. Those flagged only by more lenient criteria point to further investigation, like a more in-depth analysis of controls or subgroup analyses. Some studies clearly failed to recapitulate the exploratory results and were declared non-confirmations. This yields a framework positioning confirmatory studies as an effective screening tool for drug candidates, rather than as definitive proof of efficacy (Fig. 5). This framework potentially reduces the risk of prematurely discarding interventions that warrant further pharmacological or mechanistic validation. It moves rigorous preclinical research beyond the dichotomous decision-making largely guided by the P-value, which has been widely criticized (*31*).

**Fig. 5.**
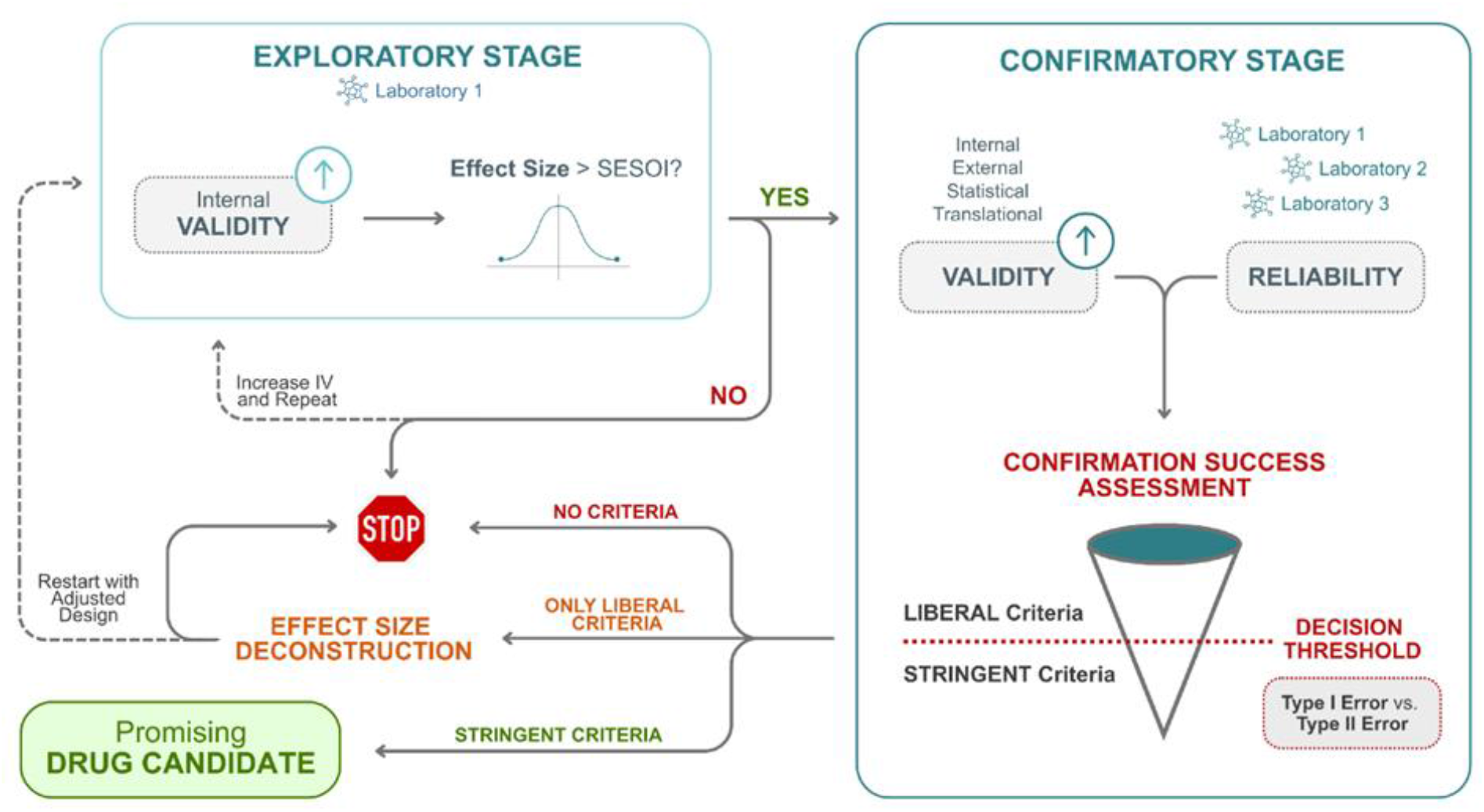
Decision-making framework for the evaluation of preclinical evidence to guide progression along the translational research pipeline. Conceptual framework for guiding translational decision-making from initial exploratory experiments to rigorous multi-lab confirmatory studies. Depending on the outcome of the confirmation assessment, promising drug candidates identified at the confirmatory stage may progress along the translational pipeline (e.g., advancing to clinical trials), undergo effect size deconstruction followed by study design adjustments, or be discontinued.

To move beyond direct replications, confirmatory studies deviated systematically from their initial exploratory counterparts. This required an in-depth knowledge of exploratory protocols and analyses, which was uniquely available in the *pCS* through the tight cooperation between the research groups and the meta-research project. All *pCS* examined here were initiated or strongly supported by investigators involved in the exploratory study. That is, researchers had sufficiently detailed knowledge of initial procedures and methods; low confirmation rates are thus unlikely to reflect major methodological shortcomings, at least for the *pCS*. This reduces concerns that a lack of technical competence, expertise, or diligence of the confirmatory laboratories explain the confirmation failure (*32*, *33*).

Based on this familiarity with the exploratory studies, researchers adjusted several experimental design characteristics to strengthen reliability and validity at the confirmatory stage. Through the implementation of strategies to mitigate risks of bias, internal validity improved in both datasets. Furthermore, using a multi-lab approach and including, for example, both sexes, strengthened the external validity and generalizability of the results. Systematically introducing differences between the exploratory and confirmatory studies, such as varying the animal strain or model induction, directly tested this generalizability (*16*, *26*). This approach yields important information about how genetic, environmental, and experimental factors interact within complex mammalian systems. Indeed, replication rates are substantially higher in genetically simpler model organisms such as *Drosophila*, where replications under standardized conditions have been comparatively robust (*34*). The improved generalizability through multi-lab studies is thus a powerful experimental tool. Improved external validity alone, however, is potentially insufficient to increase translational success (*35*). The models investigated were mostly similar to exploratory studies, leaving translational validity largely unchanged. Increased sample sizes and prespecified analysis plans contributed to the robustness and high reliability of findings. The large sample sizes reduced the chance of missing a real effect (Type II error), enabling reliable detection of even small and medium effects.

Our analysis is subject to several limitations. First, the sample size is small (19 studies), thus limiting statistical power. The two datasets and the individual studies also show great heterogeneity in methodology and underlying biology. Standardized effect sizes facilitate quantitative comparison, yet cannot fully account for differences in experimental context and biological mechanisms, so the comparability of effect sizes and validity measures may be limited. Second, the quality of the underlying data varied. For most studies in the *eCS* dataset and the exploratory *pCS* dataset, raw data were not available. We therefore had to derive effect sizes from reported summary statistics introducing measurement error and limiting the precision of effect size estimates. Third, both datasets are affected by selection bias. The *pCS* datasets is a set of studies funded by a single dedicated funding program, specifically designed as confirmatory multi-laboratory studies and with additional support from a meta-research project, limiting the generalizability of results. The *eCS* datasets consists of published studies subject to publication bias which may partly explain the higher confirmation success rate compared to the *pCS*. Fourth, comparing the exploratory and confirmatory stages is complicated by various protocol modifications or changes in animal models or induction techniques. Fifth, assessing translational validity in-depth was not feasible. A thorough evaluation requires model-specific expertise of each disease context which exceeded the scope and resources of this meta-research project. Expert elicitation was initially planned but proved unfeasible within the available timeframe. Our assessment may therefore have failed to capture differences in translational validity between exploratory and confirmatory studies. Finally, long-term outcomes such as clinical success or market approval lie further in the future. Only clinical studies will show whether preclinical confirmatory studies meaningfully improve translational development and predict clinical success better than current approaches.

Despite these constraints, this dataset provides a uniquely comprehensive look at preclinical confirmatory research covering a broad range of current multi-lab studies. The results reinforce the well-documented challenge that preclinical models often fail to translate into clinical efficacy (*36*). Although many projects did not confirm their initial exploratory findings under heightened methodological rigor, these outcomes should not be viewed as failures. Instead, because these studies explicitly incorporated adjustments to serve translational goals, their results act as a filter to prevent the advancement of putatively ineffective interventions into costly clinical trials. These findings emphasize the need to routinely fund and integrate confirmatory studies into early-stage research. Crucially, well-designed confirmatory studies yield informative evidence regardless of their outcome, thereby improving translational decisions and reducing clinical attrition.

## Materials and Methods

### The primary confirmatory studies (*pCS*) and the DECIDE project

Twelve preclinical studies, which we refer to as *primary confirmatory studies* (*pCS*), received public funding from the German Federal Ministry of Research, Technology and Space (BMFTR) (mean = €1.08 million, SD = €0.52 million; study duration: 3-5 years) to confirm a previous exploratory finding in a rigorous multi-laboratory (multi-lab) setup. The *DECIDE* project (Decision-Enabling Confirmation of Innovative Discoveries and exploratory Evidence) (*37*), a meta-research consulting team, received parallel funding and accompanied the preclinical consortia throughout their funding period (2020–2025). The DECIDE project team supported the *pCS* teams through individualized consultations, workshops, and educational webinars (see also *Supplementary Methods*) (*16*, *19*, *20*, *38–43*).

Whereas the prior exploratory experiments were conducted in a single lab, the *pCS* consisted of at least one designated *in vivo* experiment conducted across two or three independent partner labs at the national or international level (multi-lab study design). Of the twelve *pCS*, one study was unable to commence due to regulatory constraints (excluded from all analyses), another was terminated prematurely because of technical challenges (only included in study design analysis), and a third study that planned for three labs, was discontinued after two labs had produced inconclusive results (included in all analyses). Ultimately, we included eleven studies for the in-depth analysis of study design changes and ten studies for the meta-analyses.

### The extended confirmatory studies (*eCS*) dataset

We further contextualized our dataset by systematically identifying preclinical multi-lab studies reported in the literature. The analysis plan was preregistered on the OSF (*44*). We first retrieved the full texts of all studies previously identified through an existing systematic review (*45*). Using the same search string, we conducted an updated literature search twice to identify additional multi-lab studies published after the previous search date in November 2020. The study selection process is summarized in a PRISMA flow diagram (tab. S4). All records were first screened for eligibility based on title and abstract by one reviewer according to predefined inclusion and exclusion criteria (tab. S3). Preliminary inclusions were then assessed by two independent reviewers via full-text screening and were included only if studies were original, English-language preclinical research articles with full-text availability that reported primary experimental data from one multi-lab *in vivo* experiment (the same experiment conducted independently across at least two labs). Eligible studies investigated an intervention, incorporated at least one control or comparison group, and reported outcomes that were extractable for each participating lab. Discrepancies were resolved through discussion and consensus between the two reviewers or, where necessary, by consulting a third independent reviewer. For each multi-lab study, a corresponding published single-lab (“exploratory”) study investigating the same or a similar treatment and outcome measure was identified. In total, nine multi-lab confirmatory studies with corresponding single-lab exploratory studies were included (*46–61*). For each study, data were extracted from graphs, text, or supplementary materials of publications to calculate effect sizes (see also *Supplementary Methods*).

### Comparison of protocols

To collect relevant information on the experimental design changes from the exploratory to the confirmatory stage, we asked *pCS* researchers to complete structured REDCap surveys (*62*, *63*). The analysis plan was preregistered on the OSF and, together with the questionnaire content, is openly available (*64*). The survey covered four main categories: internal, external, translational, and statistical validity. Survey items included animal model, induction technique, intervention scheme, control groups, randomization and blinding procedures, and statistical analysis details. Members of 10 of the 11 *pCS* completed the surveys. For the remaining study (F), relevant information was extracted from the available publications that were shared by the study team. Initially, an additional assessment of the preregistration stage was planned, however, only 8 out of 11 *pCS* preregistered their analysis plans. Consequently, this component was not included in the analysis due to the limited sample size. Survey items were classified into different validity aspects and aggregated into preregistered and a priori-defined quantitative validity scores (tab. S11).

### Minimal internal validity (mIV) scoring

For both datasets (*pCS, eCS)*, we assessed internal validity using a preregistered, *a priori* defined mIV scoring system (minimal internal validity; mIV) based on full-text screening (tab. S12). Internal validity was evaluated across four key indicators: (i) use of controls, (ii) specification of inclusion and exclusion criteria, (iii) implementation of blinding, and (iv) use of randomization. For each indicator, one point was assigned if adequately reported. If an indicator was not reported and no justification provided, no points were assigned; if a justification for omission was provided, 0.5 points were awarded. The total mIV score was calculated as the sum of points across all four indicators. This mIV scoring system is substantially less detailed than the “*Comparison of protocols*” analysis of the *pCS* studies. However, given the limitations in reporting and extractable information in the *eCS* dataset, this simplified metric was applied consistently across datasets (*pCS*, *eCS*) to enable a quantitative comparison of methodological rigor and risk of bias.

### Meta-analyses

Effect sizes (ES) were quantified as Hedges’ *g,* with standard normal-approximation P-values (*z* = *g* / SE). Because each confirmatory study conducted experiments across multiple independent labs with differing sample sizes and conditions, lab-level data could not simply be combined into a single model. Meta-analyses instead weighted each lab’s contribution by its estimation precision. Pooled estimates were derived using fixed-effects meta-analysis, which is conventionally applied when the number of contributing labs is small (median *k* = 2) (*41*). Inter-lab heterogeneity was assessed using the I² statistic for each study, derived from fixed-effect meta-analysis models fitted using the *metagen* function (R package *meta).* Full meta-analytic results, including study-specific I² values, are reported in tabs. S6 (*pCS*) and S7 (*eCS*). Substantial heterogeneity (I² > 50%) was observed in four multi-lab studies (6, 7, E, and F), suggesting that lab-level estimates for these projects diverged more than would be expected from sampling error alone. These studies are flagged in the supplementary tables and should be interpreted with caution.

### Meta-regression

Unlike the lab-level pooling per study described above, which combines estimates of multiple labs of a common effect within each study (using a fixed-effect model), the meta-regression compares effect sizes (ES) *across* different studies, each estimating a distinct underlying effect. A random-effects model was therefore used, allowing for genuine inter-study variability in true effect size rather than assuming a single shared effect. To test whether exploratory effect size predicted the magnitude of the confirmatory (multi-lab) effect size, and whether this relationship differed between the *pCS* and *eCS* datasets, a meta-regression was fitted using the *metareg* function (R package *meta*) on a random-effects model (with studies treated as the random effect). The model was fitted as a single joint model spanning both datasets, with exploratory effect size and its interaction with dataset (*pCS* vs. *eCS*) included as moderators:

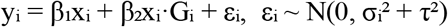

where yi and xi are the confirmatory and exploratory effect sizes for study i, Gi is an indicator equal to 1 for *pCS* and 0 for *eCS* studies, and each study’s residual variance combines its within-study sampling variance (σi²) with the between-study heterogeneity (τ², REML-estimated), so that studies are weighted inversely by their total variance. β₁ is the exploratory-to-confirmatory slope for the eCS dataset and β₂ the difference in slope between pCS and eCS. The pCS slope was derived as β₁ + β₂ with its standard error taken from the model’s variance-covariance matrix. The model was fitted without an intercept, because a treatment without an effect in the exploratory stage is not expected to show one in the confirmatory stage. Coefficients were tested with two-sided Wald z tests (α = 0.05), and the omnibus test of moderators (QM) assessed whether both slopes were zero. Residual heterogeneity was summarized by τ² and I².

### Effect size deconstruction

To attribute total effect size shrinkage to its sources, we applied a Shapley value decomposition (*65*), a method originating in cooperative game theory that distributes a total outcome fairly across contributing factors by averaging their marginal contributions across all possible orderings. We defined total shrinkage as

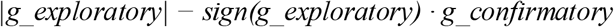

Under this formulation, a study whose confirmatory effect attenuates toward zero while remaining in the same direction as the exploratory effect, yields a shrinkage value equal to the difference in magnitude. A study whose confirmatory effect reverses direction relative to the exploratory effect yields a shrinkage value equal to the sum of both magnitudes, correctly reflecting that a directional reversal represents a more substantial departure from the original finding than simple attenuation.

Three sources were considered: control stability (change in control group mean between exploratory and confirmatory stage), treatment response attenuation (change in treated group mean), and variance inflation (change in pooled standard deviation).

For each study, we computed seven coalition effect sizes representing all possible combinations of exploratory and confirmatory component values substituted into the Hedges’ *g* formula. Confirmatory group means were estimated as sample-size-weighted averages across labs. The Shapley value for each source was computed as the weighted average of its marginal contribution across all coalitions, with weights of 2/6 for coalitions of size zero and two, and 1/6 for coalitions of size one. By construction, the three Shapley values sum exactly to total shrinkage for every project, with no residual interaction term.

A positive Shapley value indicates that the corresponding source contributed to shrinkage; a negative value indicates it decreases effect size shrinkage. The decomposition operates on raw sample-size-weighted pooled effect sizes rather than REML (restricted maximum likelihood) meta-analytic estimates; for studies with substantial between-lab heterogeneity (I² > 50%), the raw pooled confirmatory *g* may diverge from the REML estimate, and Shapley attributions for those projects should be interpreted with caution.

For each study, we applied the Distance method to detect atypical laboratories (*22*). For each lab, a statistic Dᵢ was computed as the ratio of that lab’s weighted contribution to total outcome variability relative to the overall variability across all laboratories. Dᵢ follows an F distribution under the null hypothesis of homogeneity, and labs exceeding the critical F value (α = 0.05) were flagged as atypical. The method was applied to the mean difference (control minus treated) per lab as the outcome variable.

### Z-curve analysis

To evaluate the credibility of published exploratory findings, we applied z-curve analysis (*zcurve* R package, version 2.4) to the distribution of z-scores derived from exploratory P-values. Z-curve estimates the Expected Replication Rate (ERR), defined as the average power of statistically significant studies, using an expectation-maximization mixture model. Given the minimum sample size requirement for z-curve fitting (*k* ≥ 10 significant results), exploratory studies from the *pCS* and *eCS* datasets were pooled for this analysis (n = 18 significant out of 19 total).

### Simulation framework

We simulated a two-stage preclinical trajectory comprising a single-lab exploratory experiment followed by a multi-lab (*k* = 3) confirmatory experiment. Exploratory effect sizes were sampled from empirical distributions derived from published datasets in neuroscience and metabolism studies, with robustness assessed using additional datasets from rodent fear conditioning and anxiety-related assays (*66–68*). Each confirmatory lab independently estimated the effect size, and results were pooled using a fixed-effects meta-analysis weighted by inverse variance, mirroring the analytical approach applied to the empirical confirmatory datasets.

Exploratory studies used sample sizes of n = 5, 10, 15, or 20 per group with equal allocation to treatment and control arms. Only statistically significant exploratory findings (two-sided *t*-test, P ≤ 0.05) were advanced to the confirmatory stage, reflecting common practice in preclinical research.

We modeled multiple scenarios of effect size shrinkage, ranging from 0% (optimistic scenario in which the confirmatory effect size equals the exploratory estimate) to 99% (pessimistic scenario in which no true effect is present at replication). Confirmatory sample sizes were determined via power analysis based on the observed exploratory standardized mean difference, with a maximum of n = 50 per group to reflect typical logistical constraints. Effect size attenuation was modeled using a prespecified shrinkage factor (s ∈ [0, 0.2, 0.5, 0.8, 0.99]), where confirmatory effect sizes were defined as (1 − s) times the exploratory estimate, with s = 0 indicating no shrinkage and s = 0.99 corresponding to an effect size close to zero.

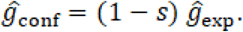

The full exploratory–confirmatory trajectory was simulated 50,000 times per shrinkage level. Confirmation success was evaluated using multiple predefined criteria (tab. S8), including significance-, direction-, interval-, and skeptical-based approaches. Performance was quantified using sensitivity (recall), false positive rate (under s = 0.99), and precision. For precision-recall analysis, low-shrinkage scenarios (s = 0 and s = 0.2) were treated as genuine confirmations (ground truth positive), and near-complete shrinkage (s = 0.99) was treated as confirmation failure (ground truth negative). Monte Carlo standard errors for all reported performance measures (replication success rates, false positive rates, precision, and recall) had a median of 0.41 % and a maximum of 3.07 %, both well below the magnitude of the differences reported between criteria.

### Confirmation assessments

For the multi-lab *pCS*, raw data provided by the *pCS* researchers was used whenever available. For the majority of exploratory studies, however, raw data was unavailable, and data points were therefore extracted from figures included in the official funding proposal or therein referenced publications, which had been shared with the DECIDE project team, using plot digitization software (*69*). Raw data of the main multi-lab *pCS* experiment were provided by the individual consortia (n = 9). One research group provided two publications resulting from their multi-lab study (F); in this case, data were extracted from these publications. Of the twelve initially funded preclinical multi-lab studies, one project could not be initiated due to regulatory constraints, and another was terminated prematurely due to technical challenges at one participating laboratory. Consequently, these two projects did not yield usable data and were excluded from the analysis.

#### Effect size estimation

Effect sizes were quantified using Hedges’ *g*, a bias-corrected standardized mean difference, to account for small sample sizes. Where primary raw data were available, group means, standard deviations, and sample sizes were calculated directly from the original datasets. Hedges’ *g* was then computed as the difference between group means divided by the pooled standard deviation, with the appropriate small-sample correction factor applied. For studies in which raw data were not available, data points were extracted from published figures using plot digitization software (*69*). Extracted values were used to approximate group means, measures of variability (standard deviations or standard errors), and sample sizes when reported.

#### Confirmation success criteria

To assess confirmatory success, defined as the extent to which multi-lab study (both *pCS* and *eCS*) confirmed their initial exploratory findings, we applied a set of criteria (tab. S8). Where raw data were available, P-values were calculated using two-sided independent-samples t-tests (or paired t-tests, where appropriate), following verification of underlying assumptions. For studies without access to raw data, P-values were either extracted directly from the original publications or reconstructed from reported summary statistics (e.g., means, standard deviations, and sample sizes) using standard parametric test formulas. The smallest effect size of interest (SESOI) was defined *a priori* based on domain-specific considerations and existing literature, representing the minimum standardized mean difference considered biologically or clinically meaningful. In the absence of established benchmarks, a conventional threshold (e.g., g = 0.2) was adopted to reflect a small effect.

### Bayesian analyses on 2 x 2 factorial comparisons

We compared minimal internal validity (mIV) scores, standardized detectable effect sizes (SDE), and experimental units (EU) between the *pCS* and *eCS* datasets using Bayesian hierarchical regression models, fitted in R with *brms* package. Each model included fixed effects for study stage, dataset, and their interaction, plus a study/stage random intercept. Because mIV scores are discrete and ordinal, they were analyzed using an ordinal cumulative probit model. SDEs, which are strictly positive, were analyzed using a lognormal model, whereas EU counts were analyzed using a Poisson model.

All models used weakly informative N (0, 1.5) priors for the fixed-effect coefficients and intercept and were fitted with four Markov chains of 2,000 iterations each. Model convergence was assessed using the potential scale reduction factor R-hat (R-hat∼1) and effective sample size (>500). Model fit was evaluated using posterior predictive checks and approximate leave-one-out cross-validation (Pareto-smoothed importance sampling with moment matching). Full posterior estimates and 95% credible intervals are reported in tabs. S9, S10 and S13.

### Additional statistical analyses

Comparisons involving two groups (RMSE, CV) were performed using the Wilcoxon rank-sum test. All statistical analyses were performed in R (version 4.5.1). The analysis code is available on <u>GitHub</u>.

## Acknowledgements

The authors used AI to assist with manuscript editing, formatting, and portions of the analysis code; all AI-assisted content was reviewed, verified, and approved by the authors. We thank everyone who was involved in the primary confirmatory studies (*pCS*) for their invaluable support, collaboration, and contributions throughout this work. We are especially grateful for the trust they placed in us by sharing raw data and for the valuable insights they provided into their study designs. A full list of contributors is available on the OSF (*64*).

## Author contributions

**Conceptualization:** CCar, MArr, NDru, PPel, SRot, UToe

**Data contribution:** ABan, ABou, ADem, AKli, AMat, ASha, BHab, BHal, FBut, FKon, GRic, HHam, HMor, ITei, ISko, JWil, KSch, KRub, LLeh, LSch, LRie, MGro, MKoc, MMei, MLöh, MTen, MVal, NBer, NDur, OMül, OPla, RSpa, SHet, SKob, SRos, TGab

**Data curation:** CCar, MArr, PPel, SRot

**Methodology**: CCar, LGam, MArr, NDru, PPel, SRot, UToe

**Analysis**: PPel, SRot

**Visualization**: PPel, SRot

**Writing - original draft:** PPel, SRot, UToe

**Writing - review & editing:** ABan, ABou, ADem, AKli, AMat, ASha, BHal, CCar, FBut, FKon, GRic, HHam, HMor, ITei, JWil, KSch, KRub, LGam, LLeh, LSch, LRie, MArr, MGro, MKoc, MMei, MLöh, MTen, NDur, NDru, OMül, OPla, PPel, RSpa, SHet, SKob, SRos, SRot, TGab, UToe

**Funding acquisition:** CCar, MArr, UToe

**Project administration**: CCar, LGam, MArr, SRot

**Supervision**: UToe

## Data, code, and materials availability

The primary confirmatory studies (pCS) raw dataset is not publicly available, as some of the underlying studies are still unpublished. Aggregated summary data sufficient to verify the manuscript’s reported results are available from the corresponding author upon request. Analysis code is available at https://doi.org/10.5281/zenodo.22871995. The protocol comparison assessment and analysis were preregistered and are publicly available (*44*, *64*). Deviations from the preregistered analysis plan are available on Zenodo (https://doi.org/10.5281/zenodo.22306511).

## List of Supplementary Materials

Materials and Methods Figs. S1 to S5

## Supplementary Methods

### The DECIDE counselling framework and educational activities

*DECIDE* (Decision-Enabling Confirmation of Innovative Discoveries and exploratory Evidence) (*37*) is a meta-research project funded by the German Federal Ministry of Research, Technology and Space (BMFTR) to accompany, support and assess the primary confirmatory study (*pCS*) groups throughout their funding period (2020–2025).

In close collaboration with the *pCS* teams, the DECIDE project team developed a counselling framework to assess and improve internal, external, and translational validity as well as reliability. The framework includes a 47-item questionnaire for systematic study assessment and the development of fit-for-purpose strategies to generate robust evidence (*16, 38*). The *pCS* received individualized consultations before study initiation and throughout the funding period, addressing topics such as preregistration, harmonization, statistical analyses, and regulatory requirements (e.g., for animal experimentation permits). All *pCS* consortia participated in an initial consultation, with several requesting follow-up meetings. In parallel, six workshops supported the development of community-driven guidelines for planning, conducting, and analyzing preclinical multi-lab studies (*16, 19, 20, 38–43*). Educational activities included four online educational seminars in 2021 and a relaunched webinar series in 2025. Recordings and slides are openly available on the Open Science Framework (OSF) (*64*).

### The *eCS* dataset: Identification of single-lab exploratory studies

We systematically identified preclinical multi-lab studies published in the literature, the extended confirmatory studies (*eCS*) dataset. We first retrieved the full texts of all studies previously identified through an existing systematic review (*45*). Using the same search string, we conducted an updated literature search twice to identify additional multi-lab studies published after the previous search date in November 2020. For each eligible multi-lab study, we further screened the publications to identify any referenced prior single-lab (“exploratory”) study that served as the basis for the multi-lab study. To be included, the exploratory study had to (i) be explicitly referenced in the multi-lab publication, (ii) be conducted in a single lab, and (iii) investigate the same or a closely related intervention, using a comparable experimental design and measuring the same primary outcome. Eight multi-lab publications with corresponding single-lab exploratory studies were included (tab. S2) (*46–61*). In the case of one publication, two independent multi-lab experiments were conducted (using different model induction techniques); these were treated as two separate multi-lab studies both linked to the same exploratory study (#4 and #5; tab. S2). In total, the *eCS* dataset comprised nine multi-lab studies with corresponding single-lab studies. In general, data from the multi-lab experiment and the corresponding single-lab experiment were extracted from graphs, text, or supplementary materials by one reviewer and cross-checked by a second independent reviewer. To calculate effect sizes (Hedges’ *g)*, means and standard deviations were extracted from each study. When only individual data points were available (e.g., presented as plots), these were extracted using plot digitization software (*69*) and subsequently aggregated into means and standard deviations. Where studies reported the standard error of the mean (SEM), these values were converted to standard deviations (SD). If multiple time points or control groups were available, the time point or control most similar to the single-lab exploratory experiment was chosen.

We aimed to select study pairs (single-lab and corresponding multi-lab experiments) with matching experimental design and primary outcomes, however, direct comparisons remain inherently limited. Several study-specific decisions were made to balance comparability with the inclusion of a sufficiently representative set of studies. In one case (#6), the single-lab and multi-lab study slightly differed in the primary outcome (infarct volume vs. lesion volume), the experimental induction technique (transient vs. permanent model induction), and measurement timing. Assessments occurred on day 1 in the exploratory study, but on day 1 (lab 1) and day 3 (lab 2) in the multi-lab study. For multi-lab study #3, we only included the data of one model (pig), while excluding two other models (rabbit, mouse) as the corresponding exploratory study was conducted in a large animal model (dog), making the pig model the most comparable. Multi-lab study #2 was excluded from variance and shrinkage analyses because its primary outcome was measured as the percentage of disease-free subjects, precluding meaningful pooling of mean values across laboratories.

### Unsupervised clustering

To assess whether *pCS* grouped by protocol similarity independently of the predefined validity classification, we performed unsupervised hierarchical clustering of relative item-level scores (score/max score, range 0–1) across all protocol items spanning the four validity domains (internal, external, statistical, and translational validity). The goal was to test whether studies and study stages (exploratory vs. confirmatory) separated based on their score profile, without imposing the four validity categories and stage as an *a priori* grouping structure. If exploratory and confirmatory stages cluster into distinct groups driven purely by the pattern of item scores, this provides independent evidence that the validity framework captures a genuine, score-driven separation between stages.

Each protocol item (e.g., blinding, randomization, control type, population criteria) constituted a row, and each study-by-stage combination (11 studies × exploratory/confirmatory stage) constituted a column, yielding a study-by-item score matrix. Hierarchical clustering (Euclidean distance, complete linkage) was applied independently to rows and columns using the *ComplexHeatmap* package in R. Validity domain annotations were displayed only as post hoc reference labels and were not used to influence clustering. Relative scores were used directly without additional scaling.

### Supplementary Figures

**fig. S1.**
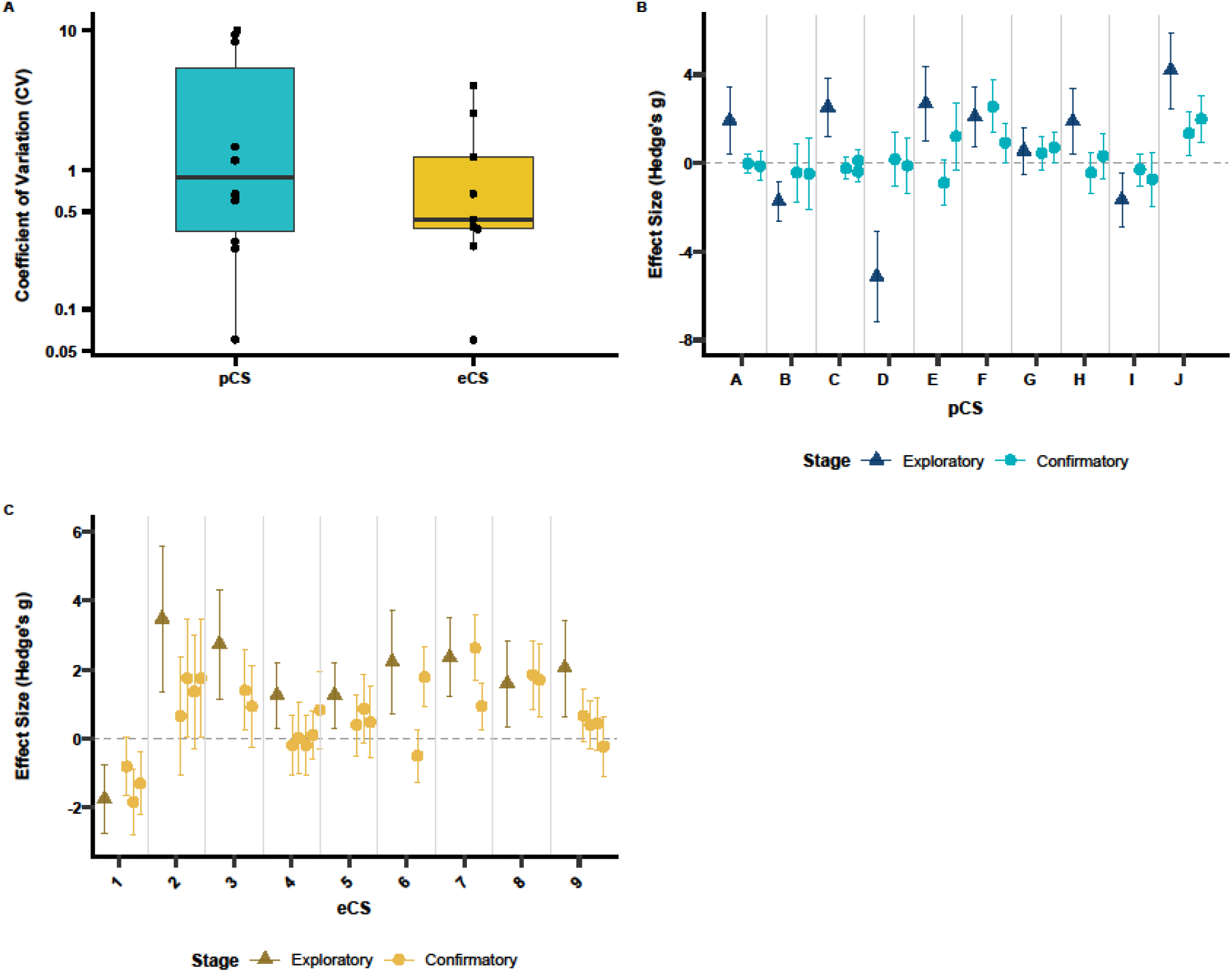
Inter-laboratory variability of the confirmatory studies **A.** Coefficient of variation (CV) of Hedges’ *g* across laboratories for *pCS* and *eCS*. CV was calculated as the standard deviation divided by the mean absolute Hedges’ *g*. Group differences were assessed using two-sided pairwise Wilcoxon rank-sum tests with Benjamini-Hochberg correction (n pCS= 10, neCS= 9, W = 54, P = 0.49). **B, C**. Change in effect size estimates from exploratory to confirmatory stage for the *pCS* (**B**) and the *eCS* (**C**). The exploratory point estimate represents the exploratory (single) laboratory, whereas confirmatory estimates represent point estimates from individual laboratories involved in one multi-laboratory study. Error bars indicate 95% confidence intervals, and the horizontal dashed line denotes the null effect.

**fig. S2.**
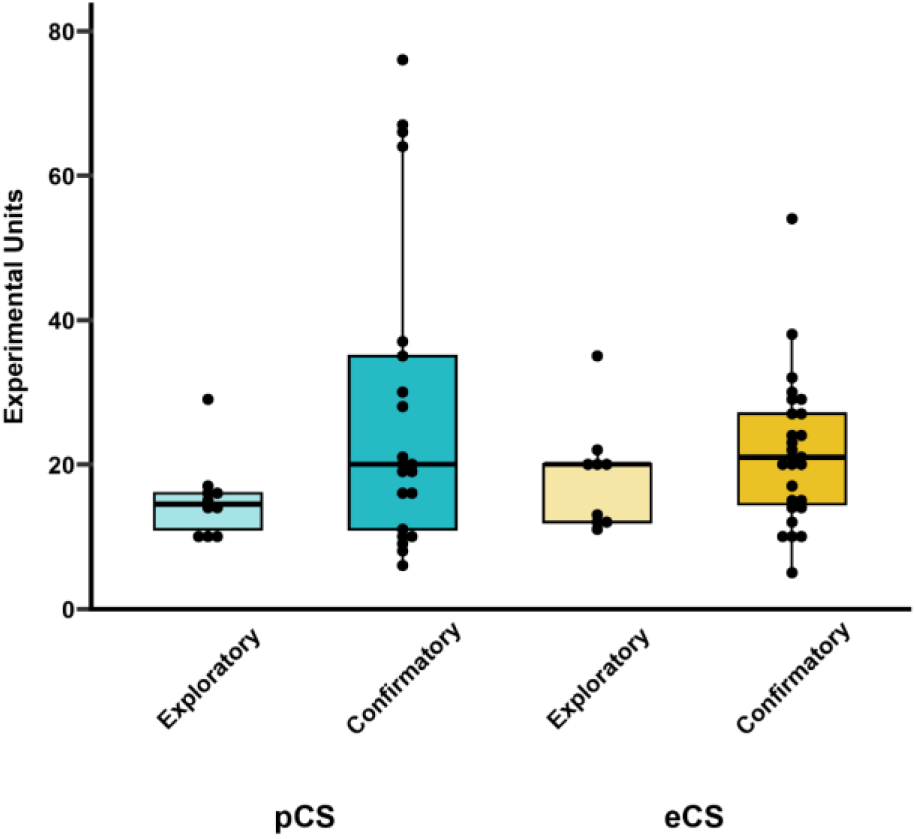
Increase in experimental units at the confirmatory stage Experimental units (EU) for primary outcomes at exploratory and confirmatory stage for *pCS* and *eCS*. The total number of units were EUpCS = 149 (exploratory) and EUpCS = 588 (confirmatory) and EUeCS = 232 (exploratory) and EUeCS = 757 (confirmatory). The total N of laboratories included in the pCS dataset was 10 (exploratory) and 21 (confirmatory). The total number of laboratories included in the *eCS* dataset was 9 (exploratory) and 31 (confirmatory). Group differences were assessed using a Bayesian hierarchical Poisson regression model with stage, dataset, and their interaction as fixed effects and a random intercept for study; 95% credible intervals excluding zero were interpreted as evidence of a credible difference.

**fig. S3.**
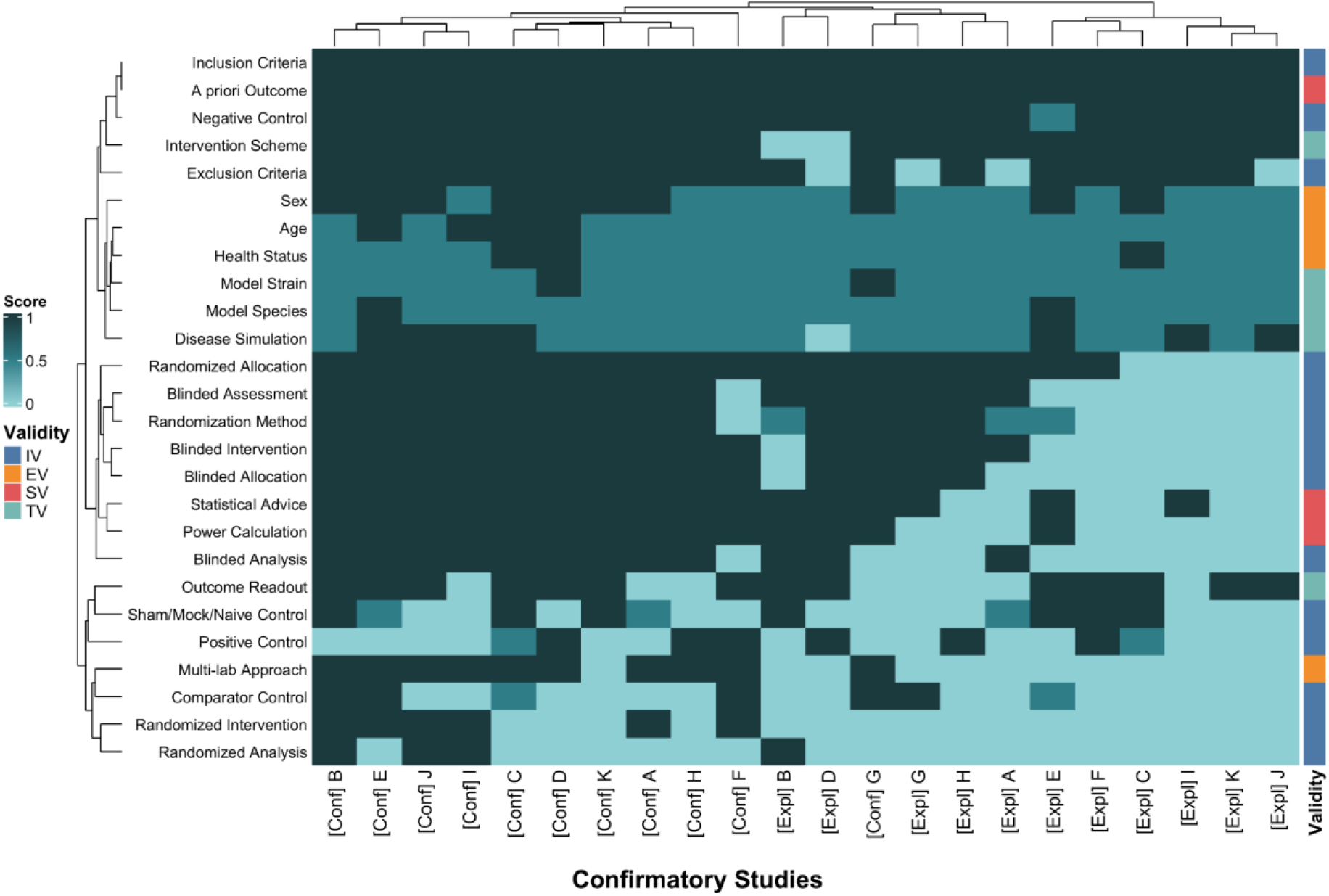
Unsupervised heatmap of the comparison of protocols Unsupervised hierarchical clustering of protocol items from different validity domains (internal, external, translational, and statistical validity) across single-lab exploratory and multi-lab confirmatory stages in the pCS (n=11). Rows represent individual methodological aspects color-coded by validity domain, and columns represent studies and study stages (exploratory, confirmatory). Teal color intensity indicates the achieved relative validity score (item score divided by its maximum, range 0 to 1). Rows and columns were clustered independently (Euclidean distance, complete linkage), and validity domain annotations were not used for clustering. Details are provided in Supplementary Methods.

**fig. S4.**
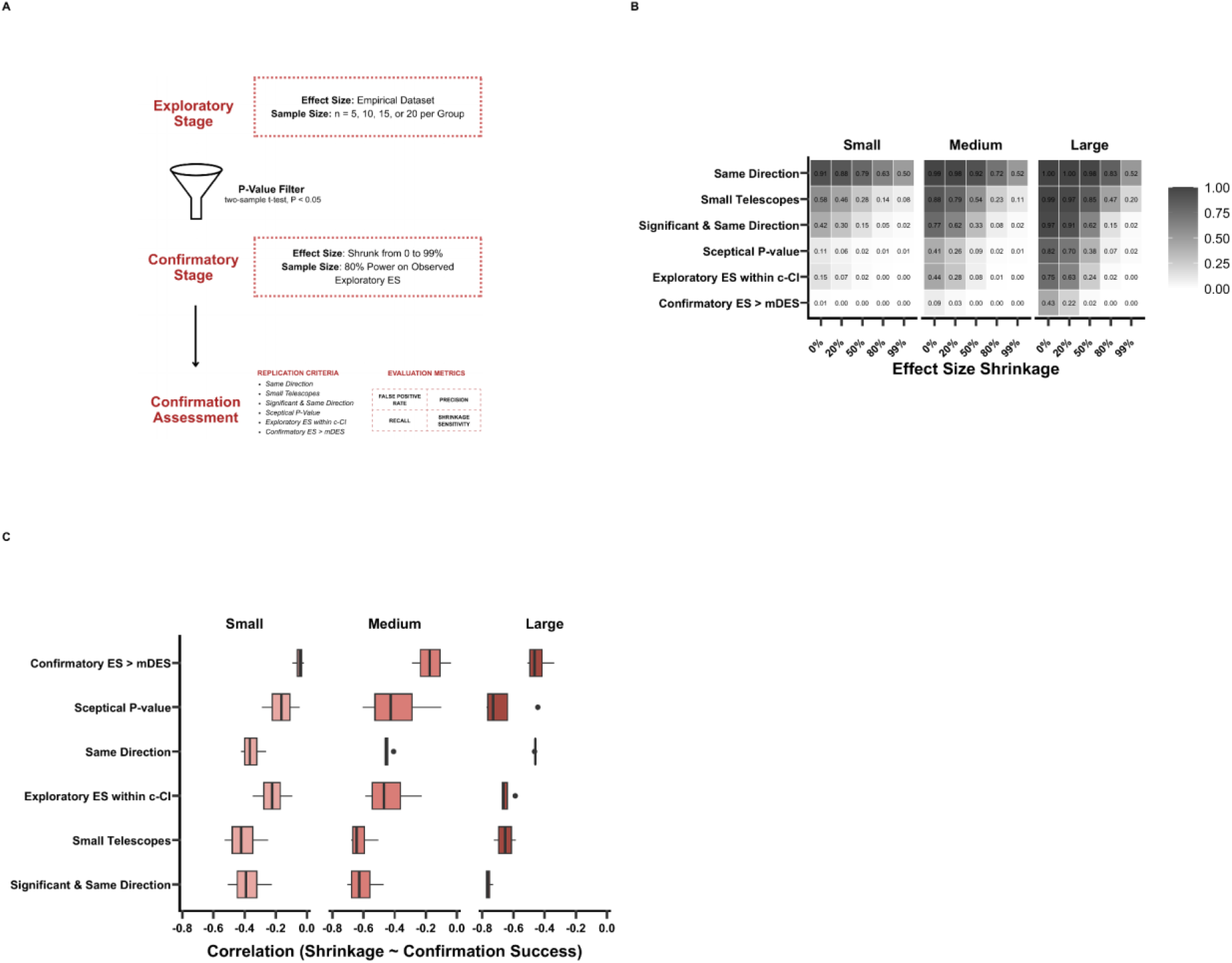
Assessment of confirmation success criteria using simulation **A.** Simulated preclinical research trajectory. Statistically significant exploratory results (two-sided *t*-test, P < 0.05; n = 5–20 per group; empirical effect size distribution) were advanced to a confirmatory stage powered at 80%, with effect size shrinkage modelled from 0 to 99%. Six confirmation criteria were evaluated on sensitivity, false positive rate, precision, and shrinkage sensitivity**. B.** Simulation-based heatmap showing confirmation success rates (color intensity) for each criterion (y-axis) across exploratory Hedges’ *g* values (x-axis) and effect size shrinkage scenarios (separate panels/gradients). Results are based on 50,000 simulated exploratory–confirmatory trajectories per condition. Exploratory n = 5–20 per group; confirmatory sample sizes were determined by power analysis and capped at 50 per group. **C.** Pearson correlation between effect size shrinkage and confirmation success plotted against exploratory Hedges’ *g* for each criterion. Each line represents a different criterion.

**fig. S5.**
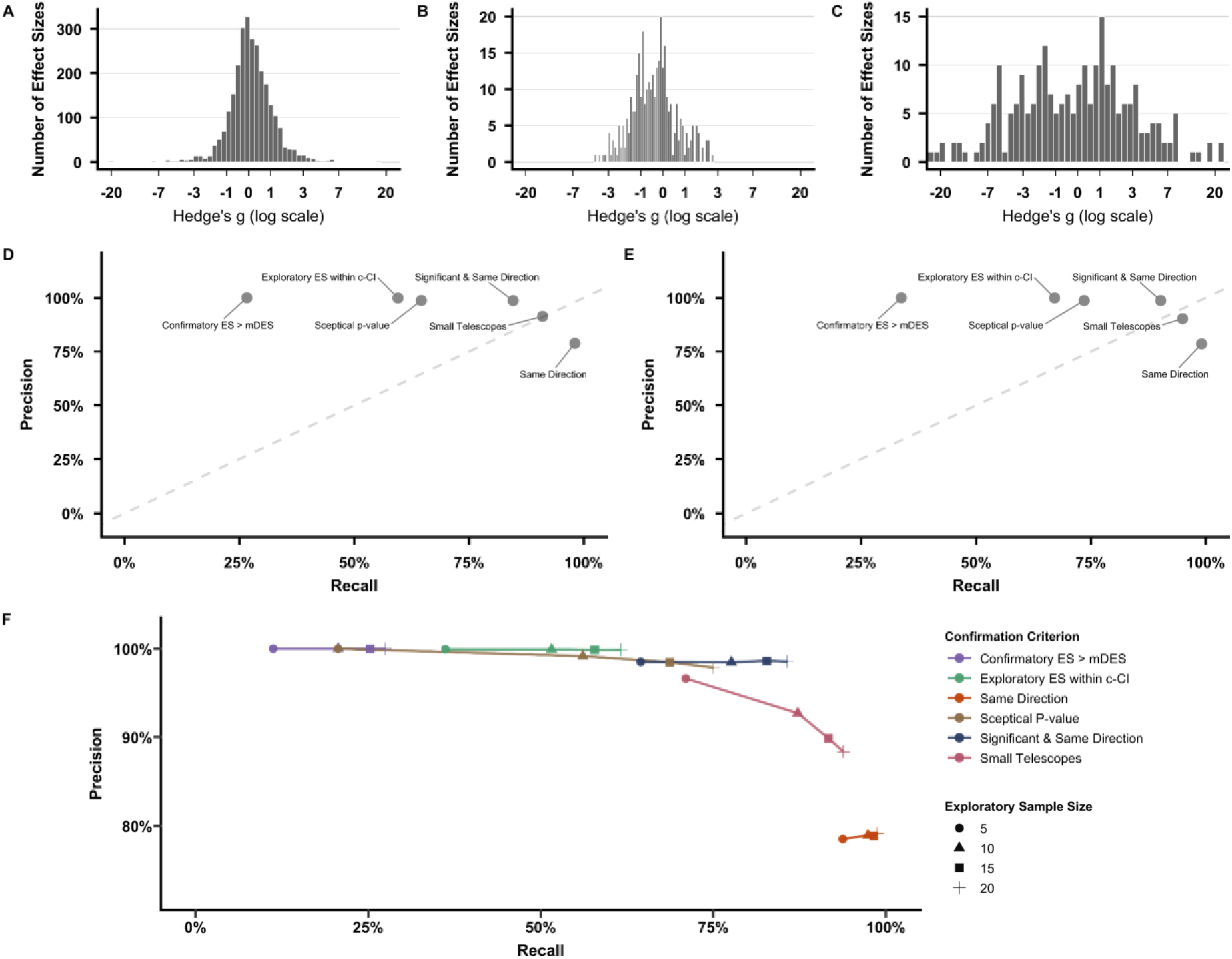
Effect size distribution from empirical datasets and sensitivity analysis of precision-recall across sample sizes. **A.** Distribution of Hedges’ *g* values derived from the primary empirical dataset (*66*). The histogram shows a right-skewed distribution (median *g* ≈ 0.8), which was used for the main simulation. **B.** Distribution of Hedges’ *g* values from a second dataset used for validation (*67*). **C.** Distribution of Hedges’ *g* values from a third dataset used for validation (*68*). **D, E.** Precision-recall curves for the datasets shown in **B** and **C**, respectively. **F.** Precision-recall curves showing the effect of exploratory sample size (n = 5, 10, 15, or 20 animals per group) on the performance of each confirmation criterion.

### Supplementary Tables

**tab. S1.** Overview of the two preclinical confirmatory dataset characteristics (*pCS*, *eCS*). Year of study conduct (*pCS*, n=11) or publication (*eCS*, n=9), research field, number and location of experimental labs per study, animal models, model induction techniques, and intervention type are shown in aggregated form for both datasets. Some studies also conducted complementary *in vitro* experiments; however, these were not included in the analyses.

|  | ***Primary confirmatory studies (pCS)***<br>(n=11) | ***Extended confirmatory studies (eCS)***<br>(n=9) |
|---|---|---|
| **Year of study conduct/publication** | 2020-2025 | 2015, 2016, 2019, 2023 |
| **Research field** | Cardiology, immunology, neurology, oncology, orthopedics, psychiatry | Cardiology, endocrinology, neurology, psychiatry |
| **Number of experimental labs** | 2-3 independent labs per study | 2-5 independent labs per study |
| **Lab locations** | Germany, Switzerland, The Netherlands | Canada, Denmark, Finland, France, Germany, Italy, Spain, Switzerland, The Netherlands, UK, USA |
| **Animal model** | Rodents (mice, rats)<br>Swine (pigs)<br>Non-human primates | Rodents (mice, rats)<br>Swine (pigs) |
| **Model induction** | Pharmacological induction<br>Substance-induced model<br>Surgical induction<br>Transgenic/knockout model<br>Vaccine-induced immune response model<br>Xenograft tumor model | Pharmacological induction<br>Surgical induction<br>Transgenic/knockout model |
| **Intervention** | Therapeutic<br>Preventive *(solely vaccination studies)* | Therapeutic |

**tab. S2.** Description of the *eCS* dataset. Published confirmatory multi-lab studies and corresponding exploratory single-lab studies, with linked publications and information on extracted experiments.

| **#** | **Exploratory *eCS*** | **Confirmatory *eCS*** |
|---|---|---|
| **1** | Modi et al. (2018)<br>Fig. 9E: Rearing Events, KO vs. rescue treatment | Arroyo-Araujo et al. (2019)<br>Fig. 1B: Rearing Events, KO vs. rescue treatment (0.63 mg/kg) |
| **2** | Ablamunits et al. (2011)<br>Fig. 1B: % Non-Diabetic [blood glucose < 250 mg/dL], placebo vs. simultaneous treatment, day 30 | Gill et al. (2016)<br>Fig. 2A: % Diabetic [blood glucose > 250 mg/dL], simultaneous placebos vs. simultaneous combinatory treatment, day 30 |
| **3** | Murry et al. (1986)<br>Fig. 3: Infarct Size [% of RaR], control vs. preconditioned | Jones et al. (2015)<br>Suppl. Online Table III: Infarct Size [% of RR]; control vs. IPC |
| **4** | Liesz et al. (2011)<br>Fig. 1B: Infarct volume [mm<sup>3</sup>] after 7 days; control vs. treatment | Llovera et al. (2015)<br>Fig. 2B: Infarct volume [mm<sup>3</sup>], control vs. treatment] |
| **5** |  | Llovera et al. (2015)<br>Fig. 2A: Infarct volume [mm<sup>3</sup>], control vs. treatment |
| **6** | Pradillo et al. (2012)<br>Fig. 1A: Infarct volume [% hemisphere], control vs. treatment; lean animals | Maysami et al. (2015)<br>Fig. 2: Lesion volume, Manchester 2012 (<sup>10 months</sup>) and Kuopio 2012, control vs. treatment |
| **7** | Garg et al. (2013)<br>Fig. 2C: Phenotypic Score, control (Mecp2-KO) vs. treatment | Powers et al. (2023)<br>Fig. 1C (NCH) and 4D (Edinburgh): Severity Score, control (Mecp2-KO) vs. treatment |
| **8** | Meloche et al. (2015)<br>Fig. 4B; systolic right ventricular pressure, RSVP [mmHg], control vs. treatment (JQ1) | Van der Feen et al. (2019)<br>Fig. 4B,E: systolic right ventricular pressure, sRVP [mmHg], control vs. intervention (RVX) |
| **9** | Gelderblom et al. (2012)<br>Fig. 5B: infarct size [mm<sup>3</sup>], control vs. treatment | Gelderblom et al. (2023)<br>Suppl. Fig. 4A: infarct size [mm<sup>3</sup>], control vs. treatment |

**tab. S3.**
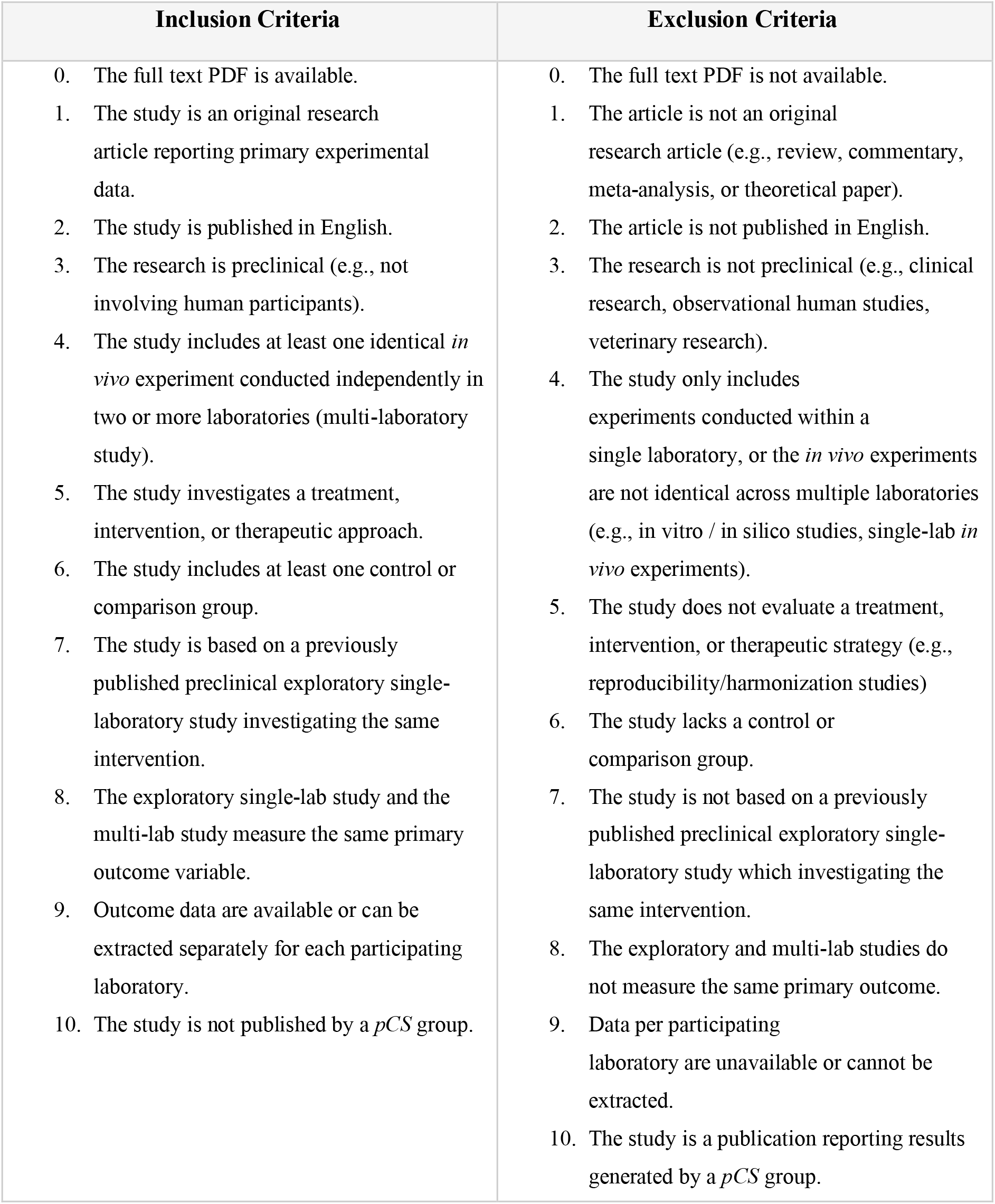
Inclusion and exclusion criteria for the *eCS* dataset. Inclusion and exclusion criteria for systematically identifying relevant preclinical multi-laboratory studies and corresponding single-laboratory exploratory studies published in the literature.

| **Inclusion Criteria** | **Exclusion Criteria** |
|---|---|
| 0. The full text PDF is available.<br>1. The study is an original research article reporting primary experimental data.<br>2. The study is published in English.<br>3. The research is preclinical (e.g., not involving human participants).<br>4. The study includes at least one identical *in vivo* experiment conducted independently in two or more laboratories (multi-laboratory study).<br>5. The study investigates a treatment, intervention, or therapeutic approach.<br>6. The study includes at least one control or comparison group.<br>7. The study is based on a previously published preclinical exploratory single-laboratory study investigating the same intervention.<br>8. The exploratory single-lab study and the multi-lab study measure the same primary outcome variable.<br>9. Outcome data are available or can be extracted separately for each participating laboratory.<br>10. The study is not published by a *pCS* group. | 0. The full text PDF is not available.<br>1. The article is not an original research article (e.g., review, commentary, meta-analysis, or theoretical paper).<br>2. The article is not published in English.<br>3. The research is not preclinical (e.g., clinical research, observational human studies, veterinary research).<br>4. The study only includes experiments conducted within a single laboratory, or the *in vivo* experiments are not identical across multiple laboratories (e.g., in vitro / in silico studies, single-lab *in vivo* experiments).<br>5. The study does not evaluate a treatment, intervention, or therapeutic strategy (e.g., reproducibility/harmonization studies)<br>6. The study lacks a control or comparison group.<br>7. The study is not based on a previously published preclinical exploratory single-laboratory study which investigating the same intervention.<br>8. The exploratory and multi-lab studies do not measure the same primary outcome.<br>9. Data per participating laboratory are unavailable or cannot be extracted.<br>10. The study is a publication reporting results generated by a *pCS* group. |

**tab. S4.**
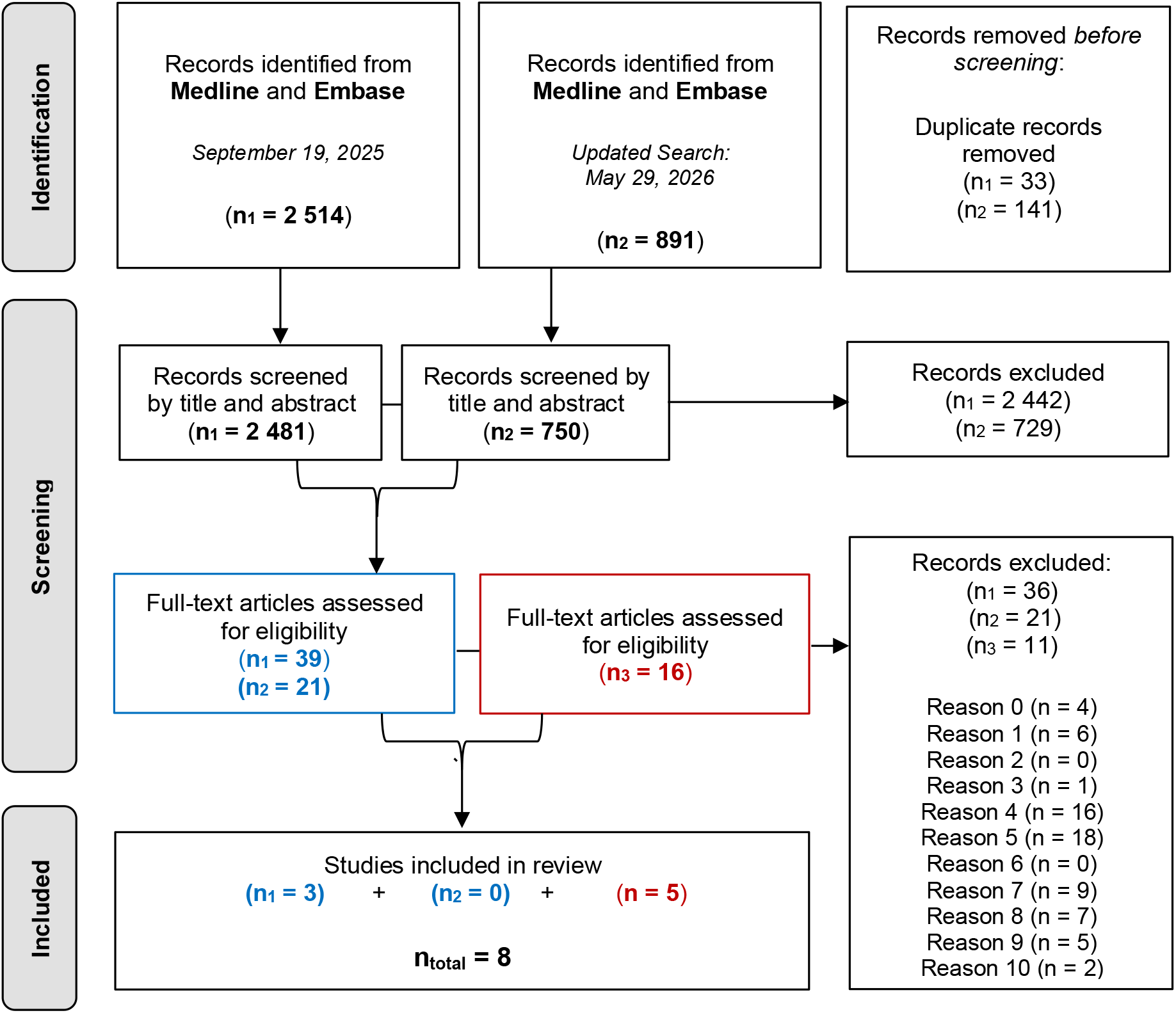
PRISMA flow diagram for the *eCS* dataset. PRISMA flow diagram of the systematic literature search used to identify relevant preclinical multi-laboratory studies. Studies shown in *red* (n3) were retrieved from a previous systematic review (45), while studies shown in *blue* (n1-2) were identified through an updated search using the same search strategy.

|  |  |  |  |
|---|---|---|---|
| Identification | Records identified from<br>**Medline** and **Embase**<br>*September 19, 2025*<br>(**n<sub>1</sub> = 2 514**) | Records identified from<br>**Medline** and **Embase**<br>*Updated Search:*<br>*May 29, 2026*<br>(**n<sub>2</sub> = 891**) | Records removed *before screening*:<br>Duplicate records removed<br>(n<sub>1</sub> = 33)<br>(n<sub>2</sub> = 141) |
| Screening | Records screened by title and abstract<br>(**n<sub>1</sub> = 2 481**) | Records screened by title and abstract<br>(**n<sub>2</sub> = 750**) | Records excluded<br>(n<sub>1</sub> = 2 442)<br>(n<sub>2</sub> = 729) |
|  | Full-text articles assessed for eligibility<br>(**n<sub>1</sub> = 39**)<br>(**n<sub>2</sub> = 21**) | Full-text articles assessed for eligibility<br>(**n<sub>3</sub> = 16**) | Records excluded:<br>(n<sub>1</sub> = 36)<br>(n<sub>2</sub> = 21)<br>(n<sub>3</sub> = 11)<br>Reason 0 (n = 4)<br>Reason 1 (n = 6)<br>Reason 2 (n = 0)<br>Reason 3 (n = 1)<br>Reason 4 (n = 16)<br>Reason 5 (n = 18)<br>Reason 6 (n = 0)<br>Reason 7 (n = 9)<br>Reason 8 (n = 7)<br>Reason 9 (n = 5)<br>Reason 10 (n = 2) |
| Included | Studies included in review<br>(**n<sub>1</sub> = 3**) + (**n<sub>2</sub> = 0**) + (**n = 5**)<br>**n<sub>total</sub> = 8** |  |  |

**tab. S5.** Meta-regression of confirmatory effect sizes on exploratory estimates. “eCS slope (β₁) “ and “Slope difference, pCS vs eCS (β₂)” represent direct model coefficients. “pCS effective slope (β₁ + β₂)” was derived post hoc as the sum of these coefficients (0.443 + (−0.304) = 0.139). Test of moderators: QM(df = 2) = 17.84, P < 0.001. Residual heterogeneity: τ² = 0.43, I² = 84.7%, QE(df = 17) = 116.61, P < 0.001. Substantial residual heterogeneity remained after accounting for exploratory effect size and its interaction with the dataset, indicating that study-specific factors contribute substantially to variability in confirmatory multi-laboratory outcomes. Abbreviations: CI, confidence interval.

| **Term** | **β** | **SE** | **z** | **P** | **95% CI** |
|---|---|---|---|---|---|
| eCS slope (β<sub>1</sub>) | 0.443 | 0.113 | 3.915 | 9.06 * e<sup>-05</sup> | [0.22, 0.66] |
| Slope difference, pCS vs eCS (β<sub>2</sub>) | -0.304 | 0.143 | -2.124 | 0.034 | [-0.58, -0.02] |
| pCS effective slope (β<sub>1</sub> + β<sub>2</sub>) | 0.139 | 0.088 | 1.587 | 0.113 | [-0.03, 0.31] |

**tab. S6.**
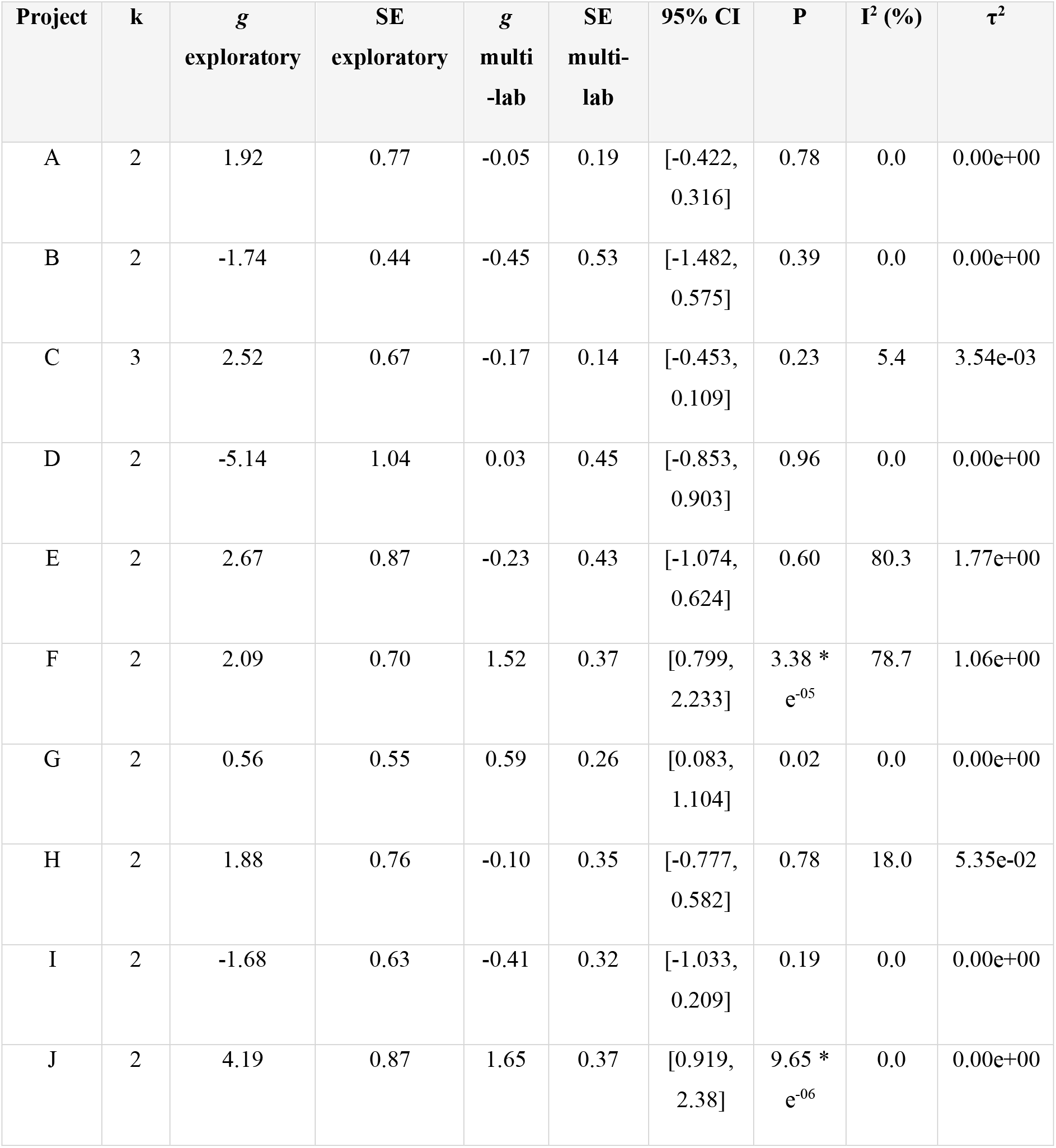
Fixed-effect meta-analysis of the *pCS* dataset. Fixed-effects meta-analysis results for the pCS (primary Confirmatory Studies) dataset, including per-project pooled effect sizes, confidence intervals, and I² heterogeneity statistics. Projects with I² > 50% are flagged with an asterisk (*) and should be interpreted with caution. Given the small number of laboratories per project (k = 2–5), I² = 0% reflects limited statistical power to detect heterogeneity rather than no heterogeneity.

**tab. S7.** Fixed-effect meta-analysis of the *eCS* dataset. Fixed-effects meta-analysis results for the eCS (extended Confirmatory Studies) dataset, including per-project pooled effect sizes, confidence intervals, and I² heterogeneity statistics. Projects with I² > 50% are flagged with an asterisk (*) and should be interpreted with caution. Given the small number of laboratories per project (k = 2–5), I² = 0% reflects limited statistical power to detect heterogeneity rather than no heterogeneity.

| **Project** | **k** | ***g* exploratory** | **SE exploratory** | ***g* multi-lab** | **SE multi-lab** | **95% CI** | **P** | **I<sup>2</sup> (%)** | **τ<sup>2</sup>** |
|---|---|---|---|---|---|---|---|---|---|
| 1 | 3 | -1.76 | 0.50 | -1.28 | 0.26 | [-1.792, -0.761] | 1.20 * e<sup>-06</sup> | 22 | 0.06 |
| 4 | 5 | 1.26 | 0.49 | 0.05 | 0.20 | [-0.349, 0.449] | 0.81 | 0 | 0 |
| 5 | 3 | 1.26 | 0.49 | 0.56 | 0.28 | [0.004, 1.116] | 0.05 | 0 | 0 |
| 6 * | 2 | 2.22 | 0.77 | 0.51 | 0.29 | [-0.064, 1.081] | 0.08 | 93.40 | 2.43 |
| 7 * | 2 | 2.35 | 0.58 | 1.51 | 0.28 | [0.954, 2.067] | 1.02 * e<sup>-07</sup> | 87.40 | 1.25 |
| 8 | 2 | 1.60 | 0.64 | 1.78 | 0.37 | [1.054, 2.507] | 1.56 * e<sup>-06</sup> | 0 | 0 |
| 9 | 4 | 2.04 | 0.71 | 0.35 | 0.20 | [-0.03, 0.732] | 0.07 | 0 | 0 |
| 2 | 4 | 3.47 | 1.09 | 1.37 | 0.43 | [0.525, 2.22] | 1.50 * e<sup>-03</sup> | 0 | 0.00 |
| 3 | 2 | 2.74 | 0.81 | 1.17 | 0.43 | [0.333. 2.001] | 0.01 | 0 | 0 |

**tab. S8.**
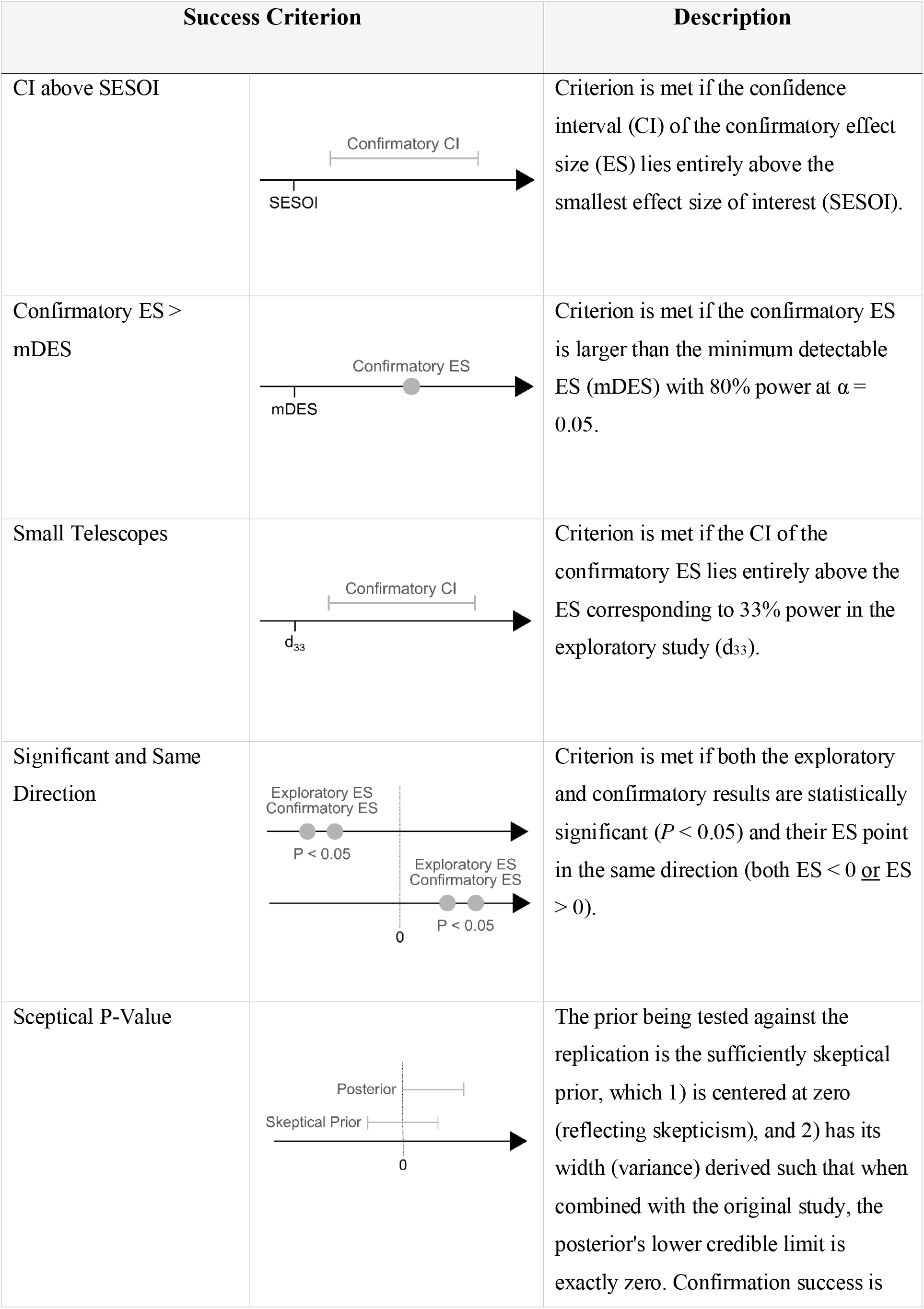
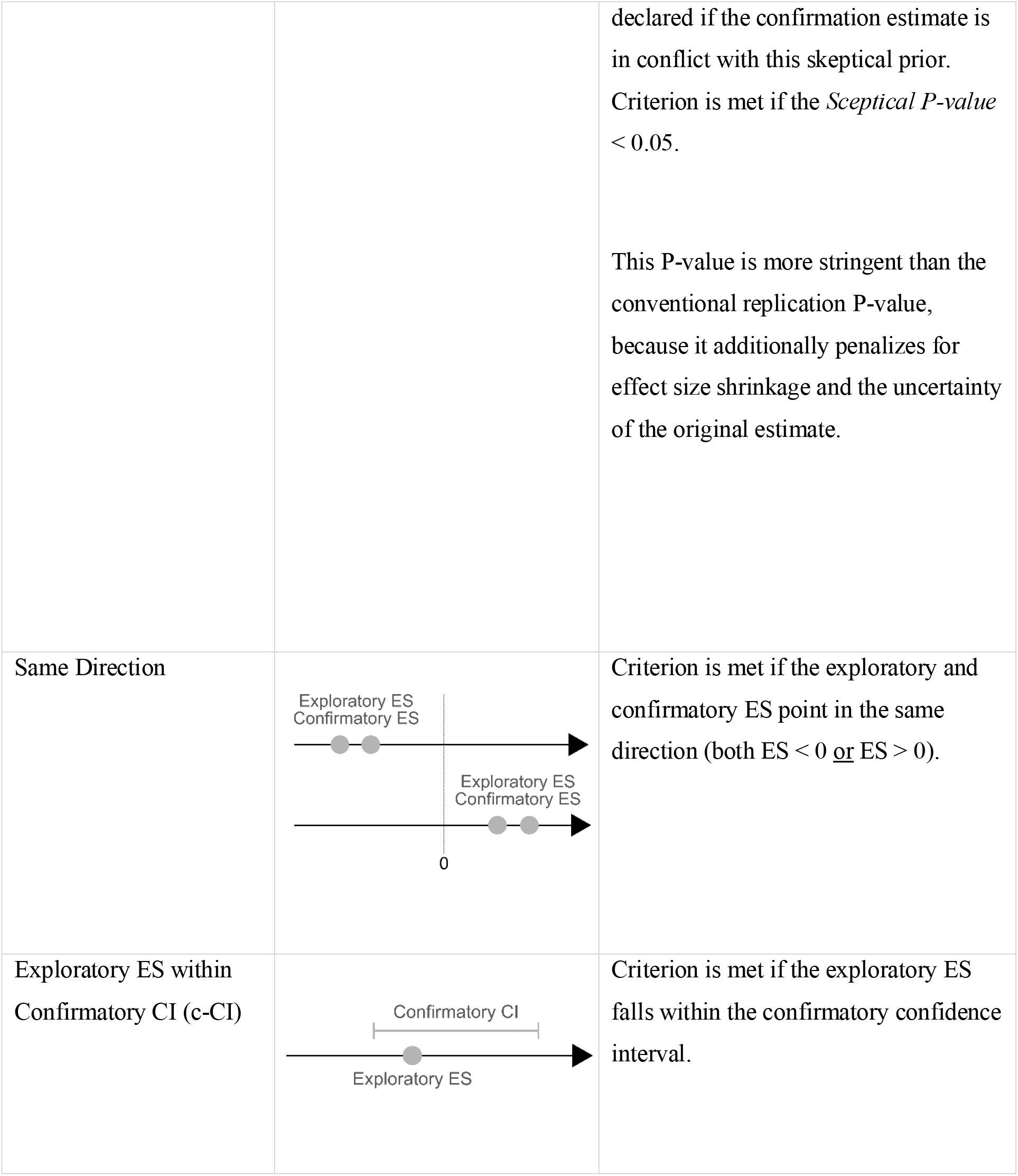
Detailed description of criteria used to assess confirmation success. . This table summarizes the statistical criteria used to evaluate confirmation, including definitions and decision rules. Abbreviations: *CI* = confidence interval, *ES* = effect size, *SESOI* = smallest effect size of interest, *mDES* = minimum detectable effect size, *d₃₃*= effect size corresponding to 33% power in the original (exploratory) study.

| Success Criterion |  | Description |
|---|---|---|
| CI above SESOI |  | Criterion is met if the confidence interval (CI) of the confirmatory effect size (ES) lies entirely above the smallest effect size of interest (SESOI). |
| Confirmatory ES > mDES |  | Criterion is met if the confirmatory ES is larger than the minimum detectable ES (mDES) with 80% power at α = 0.05. |
| Small Telescopes |  | Criterion is met if the CI of the confirmatory ES lies entirely above the ES corresponding to 33% power in the exploratory study (d<sub>33</sub>). |
| Significant and Same Direction |  | Criterion is met if both the exploratory and confirmatory results are statistically significant (*P* < 0.05) and their ES point in the same direction (both ES < 0 or ES > 0). |
| Sceptical P-Value |  | The prior being tested against the replication is the sufficiently skeptical prior, which 1) is centered at zero (reflecting skepticism), and 2) has its width (variance) derived such that when combined with the original study, the posterior's lower credible limit is exactly zero. Confirmation success is declared if the confirmation estimate is in conflict with this skeptical prior. Criterion is met if the *Sceptical P-value* < 0.05.<br>This P-value is more stringent than the conventional replication P-value, because it additionally penalizes for effect size shrinkage and the uncertainty of the original estimate. |
| Same Direction |  | Criterion is met if the exploratory and confirmatory ES point in the same direction (both ES < 0 or ES > 0). |
| Exploratory ES within Confirmatory CI (c-CI) |  | Criterion is met if the exploratory ES falls within the confirmatory confidence interval. |

**tab. S9.** Bayesian hierarchical Poisson regression of experimental units (EU). Bayesian hierarchical Poisson regression of experimental unit (EU) count. Estimates are unstandardized coefficients on the log scale with 95% credible intervals in brackets; exponentiating a coefficient yields the corresponding multiplicative effect on EU count. Reference levels: Stage = Exploratory, Dataset = *pCS*.

|  | (1) |
|---|---|
| Intercept | 2.59 |
|  | [2.24, 2.93] |
| Stage: Confirmatory (vs. Exploratory) | 0.56 |
|  | [0.39, 0.73] |
| Dataset: eCS (vs. pCS) | 0.24 |
|  | [-0.26, 0.71] |
| Stage: Confirmatory (vs. Exploratory):dataseteCS | -0.18 |
|  | [-0.44, 0.07] |
| SD (random intercept, id) | 0.45 |
|  | [0.32, 0.70] |
| Num.Obs. | 63 |
| R2 | 0.687 |
| R2 Marg. | 0.099 |
| ELPD | -270.8 |
| ELPD s.e. | 26.5 |
| LOOIC | 541.6 |
| LOOIC s.e. | 53.0 |
| WAIC | 529.7 |
| RMSE | 7.39 |

**tab. S10.** Bayesian lognormal regression of standardized detectable effect size (SDE). Bayesian lognormal regression of standardized detectable effect size (SDE). Estimates are unstandardized coefficients on the log scale with 95% credible intervals in brackets; exponentiating a coefficient yields the corresponding multiplicative effect on SDE. Reference levels: Stage = Exploratory, Dataset = *pCS*. Residual SD and the random-intercept SD are reported on the log scale.

|  | (1) |
|---|---|
| Intercept | 0.50 |
|  | [0.31, 0.67] |
| Stage: Confirmatory (vs. Exploratory) | -0.62 |
|  | [-0.88, -0.37] |
| Dataset: eCS (vs. pCS) | -0.11 |
|  | [-0.37, 0.16] |
| Stage × Dataset interaction | -0.04 |
|  | [-0.39, 0.32] |
| Residual SD (log scale) | 0.28 |
|  | [0.22, 0.37] |
| SD (random intercept, id) | 0.07 |
|  | [0.00, 0.20] |
| Num.Obs. | 38 |
| R2 | 0.661 |
| R2 Adj. | 0.505 |
| R2 Marg. | 0.648 |
| ELPD | -14.7 |
| ELPD s.e. | 5.1 |
| LOOIC | 29.5 |
| LOOIC s.e. | 10.2 |
| WAIC | 28.5 |
| RMSE | 0.27 |
| r2.adjusted.marginal | 0.541 |

**tab. S11.**
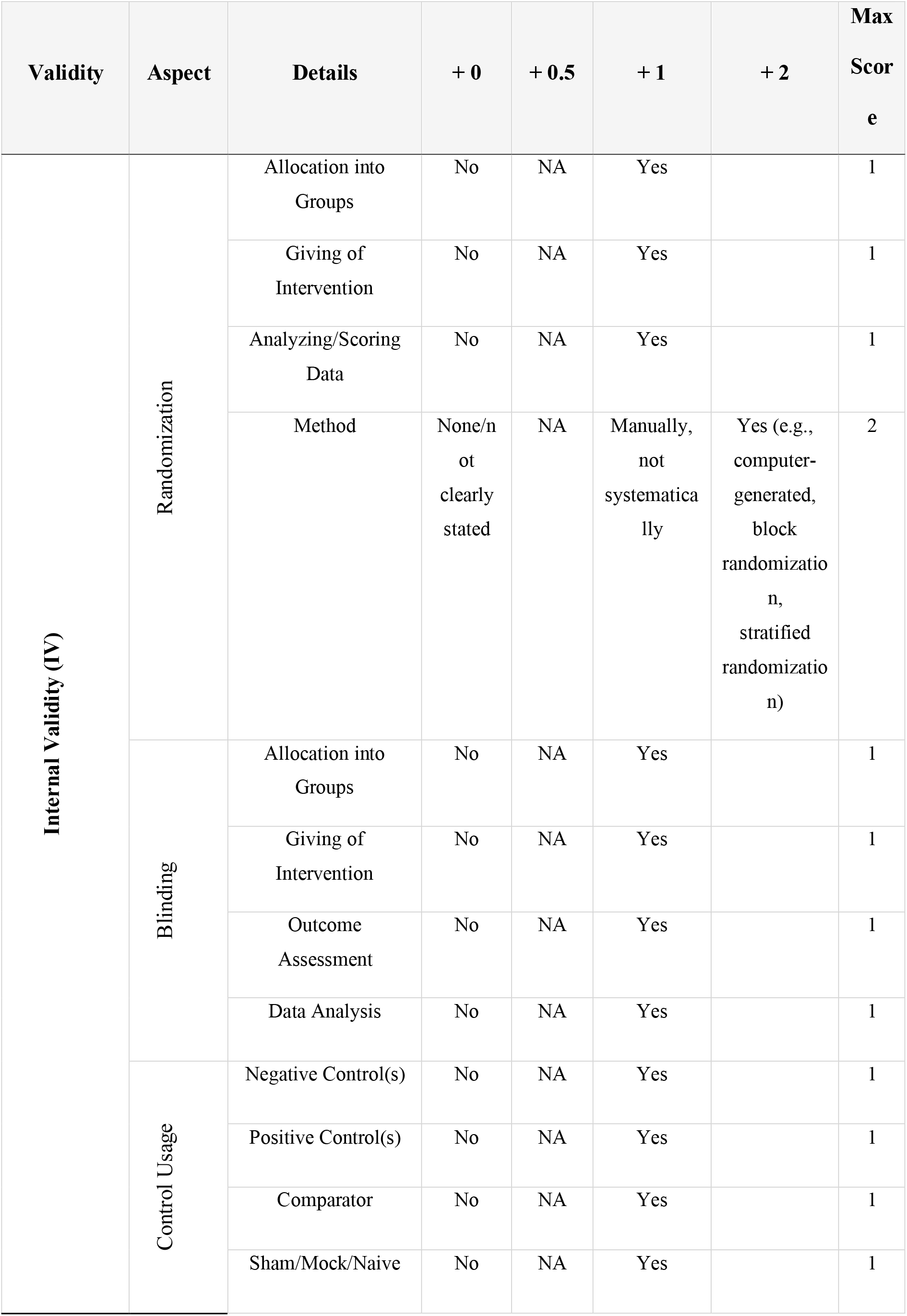
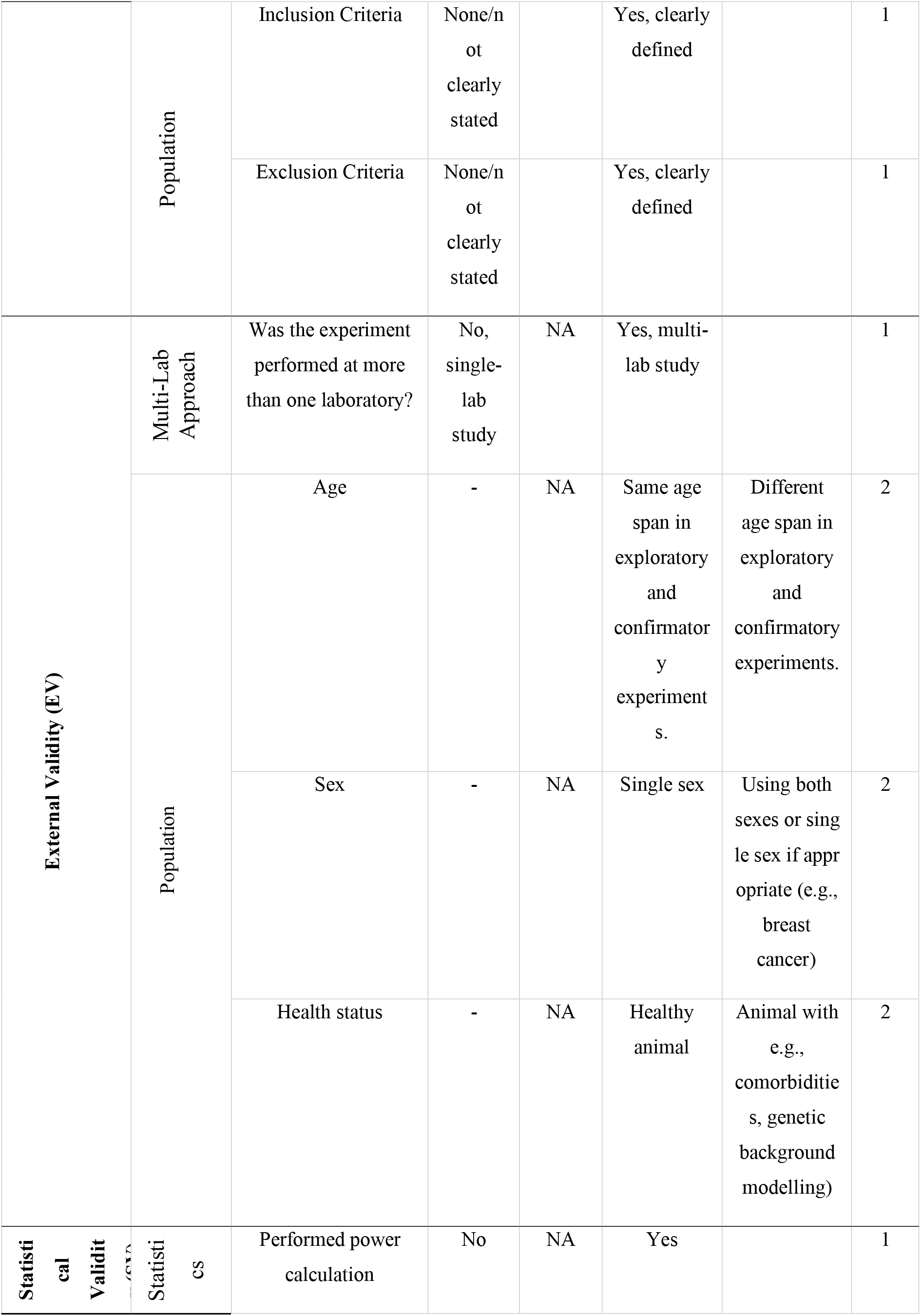
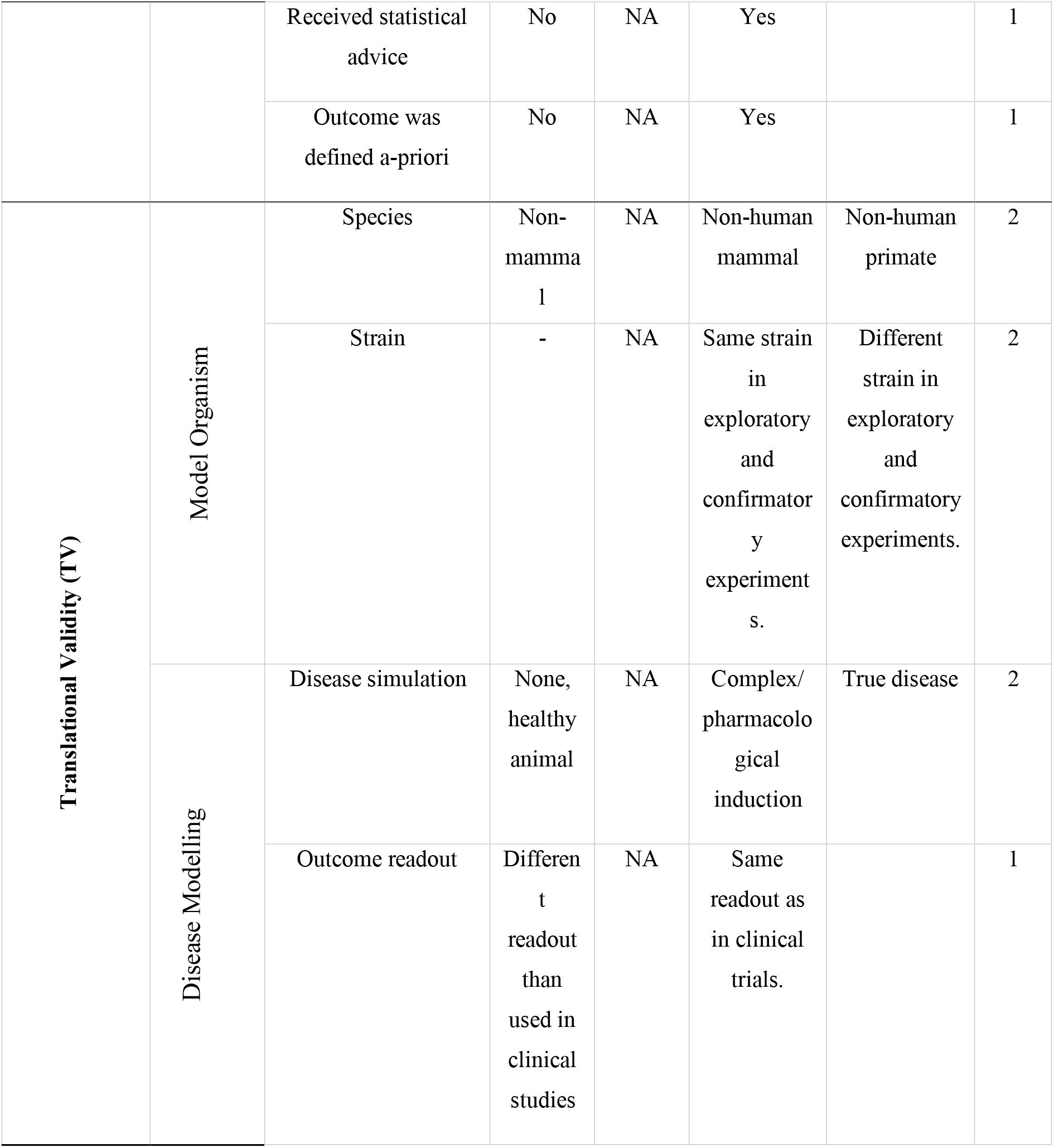
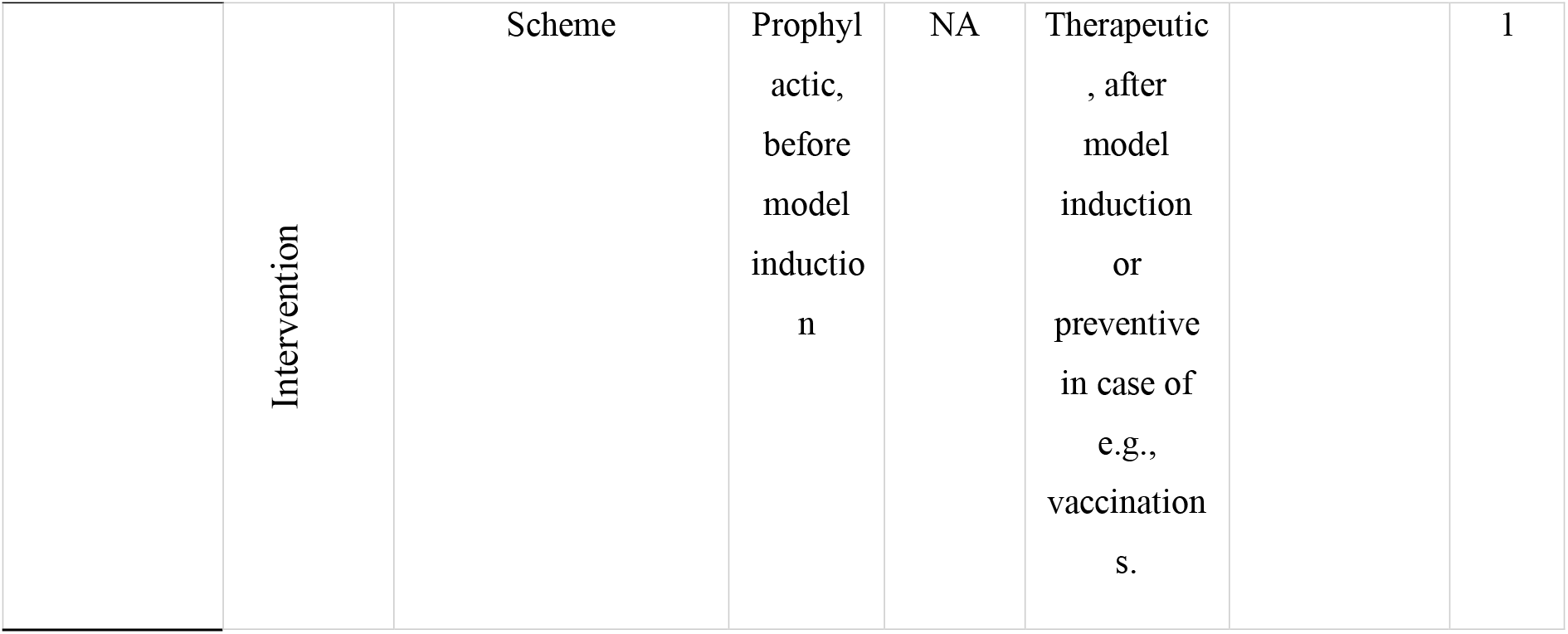
Scoring system for the *pCS* protocol comparison. . Scoring system to quantitatively assess the changes in validity (internal, external, statistical, translational validity) from exploratory to confirmatory stage for the *pCS* dataset. NA = Not applicable.

| **Validity** | **Aspect** | **Details** | **+ 0** | **+ 0.5** | **+ 1** | **+ 2** | **Max Score** |
|---|---|---|---|---|---|---|---|
| **Internal Validity (IV)** | Randomization | Allocation into Groups | No | NA | Yes |  | 1 |
|  |  | Giving of Intervention | No | NA | Yes |  | 1 |
|  |  | Analyzing/Scoring Data | No | NA | Yes |  | 1 |
|  |  | Method | None/not clearly stated | NA | Manually, not systematically | Yes (e.g., computer-generated, block randomization, stratified randomization) | 2 |
|  | Blinding | Allocation into Groups | No | NA | Yes |  | 1 |
|  |  | Giving of Intervention | No | NA | Yes |  | 1 |
|  |  | Outcome Assessment | No | NA | Yes |  | 1 |
|  |  | Data Analysis | No | NA | Yes |  | 1 |
|  | Control Usage | Negative Control(s) | No | NA | Yes |  | 1 |
|  |  | Positive Control(s) | No | NA | Yes |  | 1 |
|  |  | Comparator | No | NA | Yes |  | 1 |
|  |  | Sham/Mock/Naive | No | NA | Yes |  | 1 |
|  | Population | Inclusion Criteria | None/not clearly stated |  | Yes, clearly defined |  | 1 |
|  |  | Exclusion Criteria | None/not clearly stated |  | Yes, clearly defined |  | 1 |
| **External Validity (EV)** | Multi-Lab Approach | Was the experiment performed at more than one laboratory? | No, single-lab study | NA | Yes, multi-lab study |  | 1 |
|  | Population | Age | - | NA | Same age span in exploratory and confirmatory experiments. | Different age span in exploratory and confirmatory experiments. | 2 |
|  |  | Sex | - | NA | Single sex | Using both sexes or single sex if appropriate (e.g., breast cancer) | 2 |
|  |  | Health status | - | NA | Healthy animal | Animal with e.g., comorbidities, genetic background modelling) | 2 |
| **Statistical Validity (SV)** | Statistics | Performed power calculation | No | NA | Yes |  | 1 |
|  |  | Received statistical advice | No | NA | Yes |  | 1 |
|  |  | Outcome was defined a-priori | No | NA | Yes |  | 1 |
| **Translational Validity (TV)** | Model Organism | Species | Non-mammal | NA | Non-human mammal | Non-human primate | 2 |
|  |  | Strain | - | NA | Same strain in exploratory and confirmatory experiments. | Different strain in exploratory and confirmatory experiments. | 2 |
|  | Disease Modelling | Disease simulation | None, healthy animal | NA | Complex/pharmacological induction | True disease | 2 |
|  |  | Outcome readout | Different readout than used in clinical studies | NA | Same readout as in clinical trials. |  | 1 |
|  | Intervention | Scheme | Prophylactic, before model induction | NA | Therapeutic, after model induction or preventive in case of e.g., vaccinations. |  | 1 |

**tab. S12.**
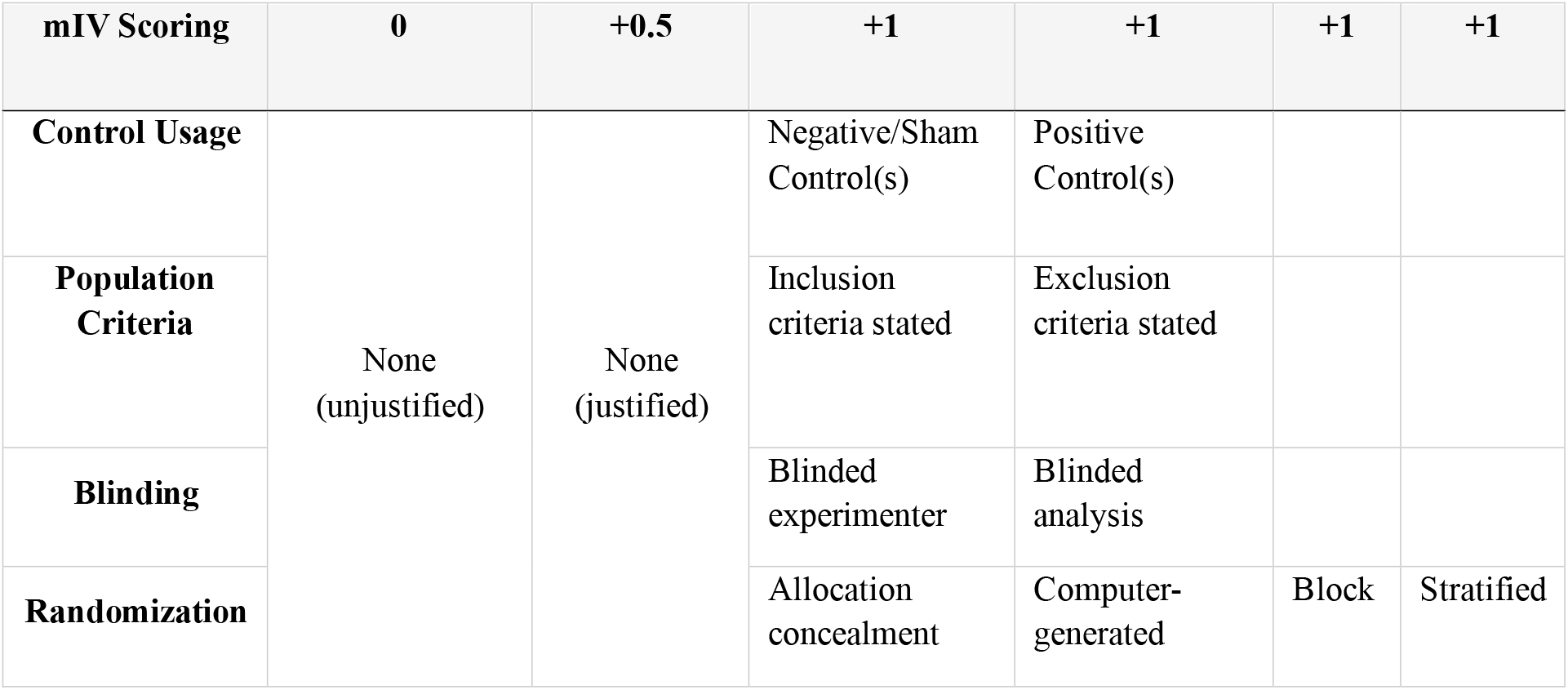
Minimal internal validity (mIV) scoring system. Scoring system to quantitatively assess the internal validity for both datasets, *pCS* and *eCS*. Block = blocked randomization; Stratified = stratified randomization by e.g., weight or sex.

| **mIV Scoring** | **0** | **+0.5** | **+1** | **+1** | **+1** | **+1** |
|---|---|---|---|---|---|---|
| **Control Usage** | None (unjustified) | None (justified) | Negative/Sham Control(s) | Positive Control(s) |  |  |
| **Population Criteria** |  |  | Inclusion criteria stated | Exclusion criteria stated |  |  |
| **Blinding** |  |  | Blinded experimenter | Blinded analysis |  |  |
| **Randomization** |  |  | Allocation concealment | Computer-generated | Block | Stratified |

**tab. S13.** Bayesian ordinal probit regression of the minimal internal validity (mIV) score. Estimates are unstandardized coefficients on the probit link scale with 95% credible intervals in brackets. Threshold parameters (1–8) denote the estimated cutpoints between adjacent mIV score categories. Reference levels: study stage = exploratory, dataset = *pCS*. Positive coefficients indicate higher predicted mIV score relative to the corresponding reference categories.

|  | (1) |
|---|---|
| Threshold 1 | -1.84 |
|  | [-2.81, -0.97] |
| Threshold 2 | -1.05 |
|  | [-1.91, -0.28] |
| Threshold 3 | -0.35 |
|  | [-1.12, 0.43] |
| Threshold 4 | -0.08 |
|  | [-0.83, 0.69] |
| Threshold 5 | 0.65 |
|  | [-0.09, 1.48] |
| Threshold 6 | 1.10 |
|  | [0.32, 2.00] |
| Threshold 7 | 1.38 |
|  | [0.57, 2.32] |
| Threshold 8 | 2.42 |
|  | [1.44, 3.60] |
| Stage: Confirmatory (vs. Exploratory) | 2.54 |
|  | [1.49, 3.67] |
| Dataset: eCS (vs. pCS) | -0.75 |
|  | [-1.78, 0.34] |
| Stage × Dataset interaction | -1.52 |
|  | [-2.91, -0.24] |
| SD (random intercept, project) | 0.58 |
|  | [0.03, 1.39] |
| Num.Obs. | 38 |
|  | (1) |
| R2 | 0.648 |
| R2 Marg. | 0.569 |
| ELPD | -71.8 |
| ELPD s.e. | 5.6 |
| LOOIC | 143.6 |
| LOOIC s.e. | 11.1 |
| WAIC | 142.1 |

